# Functional Specialization of Angular Gyrus and Precuneus Subregions for Perspective-Guided Autobiographical Memory Retrieval

**DOI:** 10.1101/2025.06.20.660447

**Authors:** Selen Küçüktaş, Peggy L. St. Jacques

## Abstract

Autobiographical memory (AM) retrieval involves goal-directed and reconstructive processes that unfold over time. A key feature of this process is the visual perspective adopted during remembering, which shapes subjective memory experience. Using fMRI, we cued participants to retrieve AMs from an own-eyes, observer, or natural perspective followed by an event probe. Our design temporally isolates preparatory (cue phase) and reconstructive (probe phase) mechanisms to identify the neural signatures of retrieval orientation, the strategic use of cues to optimize retrieval. Whole-brain and ROI analyses revealed that the angular gyrus (AG) and precuneus support perspective-guided retrieval in distinct ways. During the cue phase, PGp showed greater activity for instructed perspectives than natural retrieval, consistent with preparatory perspective selection. During the probe phase, observer-perspective retrieval elicited greater activity in AG and precuneus, supporting sustained perspective maintenance. Brain-behavior models linked PGp and 7P activity to greater vividness and perspective stability, while precuneus (7M) activity was negatively associated with emotional intensity, especially in the observer condition. These findings reveal phase- and subregion-specific contributions of posterior parietal cortex to the subjective qualities of memory. AG subregions support goal-directed perspective selection and implementation, while precuneus subregions flexibly modulate phenomenological features during memory reconstruction.

---

The retrieval of autobiographical memories (AMs) engages a dynamic unfolding of cue processing, memory search, access, and elaboration (Cabeza and St Jacques 2007). These processes are shaped by retrieval orientation, which refers to the preparatory state that optimizes access to a memory trace based on the nature of the retrieval cue (Rugg and Wilding 2000). One particularly influential cue in AM retrieval is the visual perspective from which an event is recalled (Rubin and Umanath 2015). Individuals may adopt an own-eyes perspective, visualizing the event from their original vantage point, or an observer-like perspective, viewing the scene as an external onlooker (Nigro and Neisser 1983). These perspectives not only alter the subjective experience of remembering—affecting vividness, emotion, and self-location—but also engage distinct neural mechanisms. The angular gyrus (AG) and precuneus, two subregions of the posterior parietal cortex, have been consistently implicated in perspective-taking and subjective memory reconstruction (St. Jacques 2019; 2022). Yet, it remains unclear whether these regions are involved in orienting retrieval toward a particular perspective or instead reflect neural responses that emerge once a memory has been accessed. This study examines how the brain implements visual perspective cues as retrieval orientations, and how these orientations influence the reconstruction of autobiographical memories. We advance prior research by examining subregions of the angular gyrus (PGa, PGp) and precuneus (7A, 7M, 7P), allowing for a more precise characterization of their distinct contributions during the cue and probe phases of retrieval. Specifically, we use a partial-trial design (Gilmore et al. 2018; Ollinger et al. 2001; St Jacques et al. 2013) that explicitly separates preparatory orientation from reconstruction, enabling a clean test of whether perspective cues instantiate a retrieval orientation prior to memory access.

Posterior parietal regions, particularly the angular gyrus (AG) and precuneus, play a central role in supporting the ability to adopt and update visual perspective during event memory (Faul et al. 2020; Grol et al. 2017; St Jacques et al. 2017; St. Jacques et al. 2018). The precuneus, located on the medial wall of the posterior parietal cortex, has been associated with a wide range of functions, from visuo-spatial imagery to self-related processing and episodic retrieval (Byrne et al. 2007; Cavanna and Trimble 2006). Functional neuroimaging studies indicate that this region is especially involved when people shift their visual perspective from their natural viewpoint (St Jacques et al. 2017; St. Jacques et al. 2018), and temporary lesions here disrupts scene constructive processes that support egocentric based memory retrieval (Hebscher et al. 2020). In contrast, the angular gyrus, on the lateral surface, has been implicated in the integration of multimodal event features and self-location within memories (Bonnici et al. 2016; Humphreys et al. 2021). Rather than supporting perceptual transformations per se, AG appears to contribute more broadly to reconstructing events in novel ways (Faul et al. 2020; St. Jacques et al. 2018), including representing bodily aspects of visual perspective (Iriye and St. Jacques 2024). Together, these regions form part of a posterior medial network that facilitates dynamic transformations of memory content across time, space, and viewpoint (Ritchey and Cooper 2020).

The precuneus and angular gyrus are anatomically expansive and functionally heterogeneous, making it difficult to isolate their specific contributions to visual perspective during different stages of memory retrieval. For instance, the precuneus comprises three major cytoarchitectonic subregions (7A, 7M, and 7P), which are topographically organized and functionally distinct (Scheperjans et al. 2007). Area 7A, located in the anterior portion of the precuneus, has been associated with self-centered mental imagery and attentional processes (Cavanna and Trimble 2006; Scheperjans et al. 2007). Functionally, the anterior precuneus has connectivity with the superior parietal cortex, paracentral lobule, and motor cortex, suggesting a sensorimotor role (Jitsuishi and Yamaguchi 2023; Margulies et al. 2009). Area 7M, situated in the posterior and ventral portion of the precuneus, is part of the default mode network associated with episodic memory retrieval and self-related processing through connections to medial prefrontal cortex, medial temporal lobe, and posterior cingulate cortex (Jitsuishi and Yamaguchi 2021; Margulies et al. 2009). Notably, area 7M has been linked to the emotional modulation of perspective shifts during memory retrieval, contributing to reductions in emotional intensity when adopting an observer viewpoint (St Jacques et al. 2017). Area 7P, located in the posterior portion of the precuneus, is thought to contribute to visual abilities through functional connectivity with adjacent visual cortical regions (Margulies et al. 2009). Similarly, the AG can be subdivided into PGa (anterior) and PGp (posterior) based on cytoarchitectonic and functional criteria (Caspers et al. 2008; Caspers et al. 2006). PGa has been associated with semantic integration and conceptual processing, potentially supporting the alignment of retrieval goals with perspective cues (Binder and Desai 2011; Seghier 2013), while PGp is more closely linked to episodic memory and subjective recollection (Bonnici et al. 2016; Humphreys et al. 2021). Disruption to AG, particularly PGp, impairs the ability to recall events from an own-eyes perspective and reduces episodic detail (Berryhill et al. 2007; Bonnici et al. 2018; Yazar et al. 2017). Together, these findings suggest that adopting a visual perspective involves reconstructing spatial and sensory details through flexible, subregion-specific contributions of the posterior parietal cortex. To enhance anatomical precision and reproducibility, we leverage cytoarchitectonically defined ROIs within AG (PGa, PGp) and precuneus (7A, 7M, 7P) during preparatory (cue) and reconstructive (probe) phases of autobiographical memory retrieval.

In prior studies, visual perspective cues were often presented simultaneously with the memory probe (e.g., an event title), likely triggering ecphory, the interaction between the cue and stored memory trace, at the time of presentation. For example, in Iriye and St. Jacques (2020), participants received a memory title alongside a visual perspective cue and were asked to search for and elaborate on a related memory. Although the study provided important evidence that visual perspective modulates early functional connectivity between the hippocampus and a posterior medial network it could not disentangle cue-related processes from memory retrieval due to their temporal overlap. The co-presentation of cues and memory probes makes it difficult to isolate neural responses to the perspective cue itself, confounding cue processing with memory access. To address this gap, we implemented a cue-probe design that temporally separates the visual perspective cue from the autobiographical memory probe (Gilmore et al. 2018; Ollinger et al. 2001; St Jacques et al. 2013). This enables us to isolate a pre-retrieval phase in which participants adopt a perspective before retrieving the memory. By separating the cue from the memory probe, our design allows us to investigate visual perspective as a form of retrieval orientation, the top-down specification of how a memory should be accessed, based on the nature of the cue (Rugg and Wilding 2000). While retrieval orientation has traditionally been studied using manipulations of cue modality or semantic content (Dobbins and Wagner 2005; Herron and Rugg 2003; Morcom and Rugg 2012), it has rarely been applied to AM. A notable exception is a study by Gurguryan and Sheldon (2019), who found that orienting AM retrieval toward contextual versus conceptual information produced distinct neural signatures in posterior parietal, medial temporal, and frontal regions. These findings suggest that retrieval orientation can shape the neural processes involved in constructing AMs. By testing perspective as a retrieval orientation, we refine accounts of posterior medial network function, specifying when and how posterior parietal cortex subregions bias the format of recollection rather than simply its magnitude.

In the current study, we examined whether visual perspective cues (own-eyes or observer) can function as retrieval orientations that shape subsequent memory reconstruction. During fMRI scanning, participants were cued to retrieve AMs from one of three conditions: own-eyes perspective, observer perspective, or a natural/unconstrained perspective (Retrieve). On some trials, participants were presented with the title of a memory they had generated one week prior to the scanning session, whereas other trials preceded directly to fixations. This fMRI design enabled us to compare neural and behavioral responses during both the cue and probe phases (Gilmore et al. 2018; Ollinger et al. 2001; St Jacques et al. 2013). We predicted that the angular gyrus and precuneus would show increased activation during the cue phase for perspective-guided (own eyes or observer) versus natural retrieval cues, consistent with the adoption of a retrieval orientation. Importantly, we examined anatomically defined subregions of within the AG (PGa and PGp) and precuneus (7A, 7M, and 7P) to identify whether distinct subregions differentially support preparatory versus reconstructive retrieval processes. We tested the hypothesis that anatomically distinct subregions of the AG and precuneus would differentially contribute to preparatory (cue) and reconstructive (probe) phases of AM retrieval, particularly in relation to visual perspective. Specifically, we expected that anterior subregions (PGa, 7A) would be engaged during the cue phase, consistent with preparatory processes such as attentional orienting and goal specification (Margulies et al. 2009; Seghier 2013), while posterior subregions (PGp, 7P) would show increased activity during the probe phase, supporting reconstruction of episodic content and perspective maintenance (Bonnici et al. 2016). We also predicted distinct brain-behavior relationships across regions: PGp and 7P recruitment would positively predict vividness, while 7M activation would be associated with reduced emotional intensity during observer-perspective retrieval, reflecting its role in emotional modulation of perspective shifts (St Jacques et al. 2017).

## Materials and Methods

### Participants

Thirty-one right-handed, healthy young adults, aged 18 to 30, were recruited for the study. The recruitment was restricted to individuals with no history of neurological or psychiatric disorders, who are not taking any medications affecting mood or cognition, and who have normal or corrected-to-normal vision. Data from one participant was excluded due to providing missing responses to 18% of the ratings during fMRI scanning. Thus, the final sample included 30 participants (20 females, *M*_age_ = 23.00, *SD* = 3.48). The sample size was determined based on a previous study investigating neural recruitment while remembering AMs from own eyes and observer-like perspectives (St. Jacques et al., 2017; 2018). Informed consent was obtained from all participants for being included in the study. Written consent was obtained. The study was approved by the University of Alberta Research Ethics Board (Pro00122284).

### Procedure

The study involved two sessions, separated by one week. Session 1 (AM generation) took place in the lab. Participants were asked to recall 90 specific AMs that occurred five years ago or older. We asked participants to remember remote events as we aimed to capture variability in the initial visual perspective adopted (Rice and Rubin 2009). For each event, participants were asked to provide a brief title and the year when the event occurred, and rate the event in vividness, emotional intensity, positive valence, own eyes, and observer-like perspectives on a 5-point scale (1=low; 5=high). Among 90 events recalled in Session 1, we selected 72 to randomly assign to Own Eyes, Observer, or Retrieve conditions in Session 2 (see below for details), such that the initial event ratings and remoteness did not differ across conditions.

Session 2 (fMRI scanning) took place approximately one week later (*M* = 7.35 days, *SD* = 1.14). Each trial during the scanning consisted of a cue and a probe phase. In the cue phase (4 s), participants were presented with a cue, indicating that they should prepare to adopt an “Own Eyes”, “Observer” perspective, or to simply “Retrieve.” In the Own Eyes condition, participants were asked to remember the event from an own eyes perspective by visualizing it through their own eyes. In the Observer condition, participants were asked to remember the event from an observer-like perspective, as if they could see themselves and their surroundings. Finally, in the Retrieve condition, participants were instructed to remember the same event again as it is naturally experienced. The retrieve condition served as a control condition by requiring participants to prepare for memory retrieval without a specific retrieval orientation for visual perspective. Importantly, although no memory title was presented during the cue phase, participants were explicitly instructed to prepare for memory retrieval by adopting the visual perspective indicated (Own Eyes or Observer). This type of cue-driven preparation is consistent with the broader literature on retrieval orientation, which shows that participants can proactively configure their mental state in response to goal-relevant cues even before memory access occurs (Dobbins and Wagner 2005; Herron and Rugg 2003; Morcom and Rugg 2012). In our design, this meant preparing to remember from a specific visual vantage point (Own Eyes or Observer) or getting ready to retrieve a specific event (Retrieve), even if the memory content had not yet been specified. Such preparatory states are thought to bias subsequent memory search and reconstruction.

The cue phase was followed by the probe phase (9 s), in which the event titles that participants generated in Session 1 were presented. Participants were asked to recall the event by adopting the visual perspective presented in the cue phase (or retrieve it as is). Then, they rated each event in emotional intensity, vividness, and perspective maintenance (i.e., how well they maintained the adopted visual perspective in that specific trial) on a 5-point scale (1=low to 5=high; each rating was presented for 3 seconds). The order of the retrieval cues was counterbalanced for each participant, and each event was recalled once. Before scanning, participants were given a training task with event titles that were not shown during scanning to familiarize them with the instructions. The fMRI scanning consisted of six functional runs, each with a total of 18 trials. Of these, 12 were full trials (i.e., cue phase followed by probe phase and ratings), and 6 were partial trials (i.e., cue phase only).

To dissociate preparatory (cue) and reconstructive (probe) phases of autobiographical memory retrieval, we implemented a compound event-related design using both full and partial trials (Ollinger et al., 2001; St. Jacques et al., 2013; Gilmore et al., 2018). Cue and probe regressors were modeled as boxcar functions with durations of 4 seconds and 9 seconds respectively, convolved with a canonical hemodynamic response function (HRF). Cue-related regressors included both full and partial trials, while probe-related regressors included full trials only. Although the onset of the probe always followed the cue after a fixed interval, we included a variable intertrial interval (ITI; 2–10 s, exponentially distributed) following both full and partial trials to reduce collinearity between events across the run. Critically, modeling cue and probe phases with appropriate durations allowed us to account for sustained cognitive engagement, and the use of partial trials enabled us to estimate non-overlapping BOLD responses despite temporal proximity. This approach has been successfully validated in previous studies (e.g., St. Jacques et al., 2013) and permits reliable estimation of cue- and probe-related activity even without a finite impulse response (FIR) model. Although FIR models are often used for dissociating closely spaced events (Ollinger et al., 2001; Gilmore et al., 2018), our partial-trial design with variable ITIs permitted reliable estimation using canonical HRF regressors. Importantly, supplementary FIR analyses confirmed that the cue and probe phases elicited distinct time courses (Supplementary Figure 1), validating the separability of our main GLM estimates. Additionally, collinearity diagnostics indicated acceptable levels of regressor independence based on variance inflation factors (*M* = 2.16, *SD* = 0.21). Taken together, these modeling choices support the validity of our approach to isolating cue and probe responses.

Trials were separated by an active baseline involving left/right decisions to eliminate the resting state activity between trials and better contrast the recruitment in cue and probe phases (Stark and Squire 2001), which were equally spaced across a variable length (2 to 10 seconds) and distributed exponentially such that shorter inter-trial intervals occurred more frequently than longer ones (see Figure 1).

**Figure 1.**
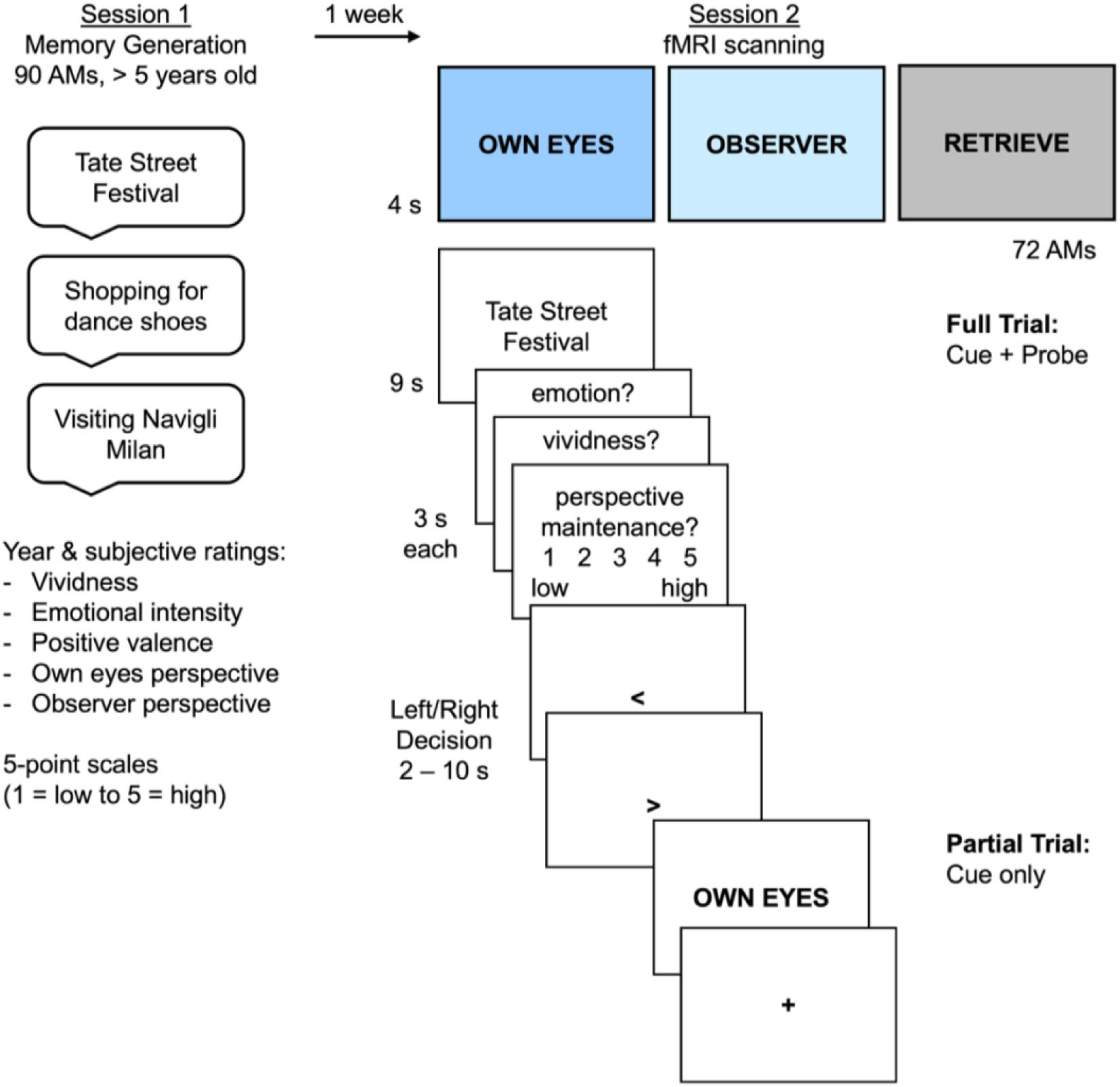
Study design. In Session 1, participants generated 90 AMs that occurred 5 years ago or older and provided a brief title, date, and subjective ratings for each event. Seventy-two of the events recalled in Session 1 were selected for Session 2 and randomly assigned to Own Eyes, Observer, and Retrieve conditions, ensuring that event remoteness and subjective ratings matched across conditions. In Session 2, participants underwent fMRI scanning while being presented with full or partial trials. In full trials, they were presented with a cue (Own Eyes, Observer, Retrieve) preparing them to retrieve events presented in the probe (AM title) by adopting an own eyes perspective, observer-like perspective, or to retrieve them as naturally experienced, respectively. Following each retrieval, participants gave subjective ratings in emotional intensity, vividness, and perspective maintenance. In partial trials, they were presented with the cue only, without probes.

### fMRI Data Acquisition and Pre-processing

Functional and structural images were acquired on a 3T Siemens Magnetom Prisma Scanner at the Peter S. Allen MR Research Centre at the University of Alberta Hospital. A desktop computer running PsychoPy (2022.2.2) software (Peirce et al. 2019) controlled stimulus display via a projector screen at the head of the scanner. Participants viewed the screen through a mirror attached to the head coil of the scanner. Cushions were used to minimize head and body movements, and earplugs were provided to attenuate scanner noise. Participants were provided a 5-button box under their right hand to provide responses.

Anatomical images were acquired using a high-resolution three-dimensional magnetization-prepared rapid gradient echo sequence (MPRAGE; 208 sagittal slices per slab, echo time [TE] = 2.37 ms, repetition time [TR] = 1800 ms, inversion time [TI] = 900 ms, flip angle = 8 degrees, voxel size = 0.9 x 0.9 x 0.9 mm). Functional images were acquired using T2* gradient echo, echoplanar imaging (EPI) sequence sensitive to BOLD contrast (TE = 30 ms, TR = 2000 ms, TI = 900 ms, flip angle = 80 degrees, voxel size = 2.2 x 2.2 x 2.2 mm). Whole-brain coverage was obtained with 64 contiguous slices acquired in the coronal orientation.

Imaging data were preprocessed and statistically analyzed using SPM12 (Wellcome Department of Imaging Neuroscience, London, UK) using standard methods. The data were preprocessed to remove noise and artifacts, which included slice-timing correction, realignment, co-registration to align anatomical and functional images for each participant, segmentation, normalization to the Montreal Neurological Institute (MNI) template, and smoothing using a Gaussian kernel (5 mm full-width at half maximum [FWHM]).

### Behavioral Data Analysis

Jamovi (2.3.28) was used for all statistical analyses. Unless otherwise stated, all statistical analyses were run using a repeated-measures ANOVA with Retrieval Cue (Own Eyes, Observer, Retrieve) as the within-subject factor. Separate analyses were conducted for emotional intensity, vividness, and perspective maintenance ratings. Greenhouse-Geisser correction was reported when the assumption of sphericity was violated. Holm-Bonferroni adjustments were reported to control for pairwise comparisons.

### fMRI Analysis

#### Whole-Brain Analysis

A general linear model (GLM) approach was used to analyze the fMRI data and included regressors at the onset of each cue and probe. Cue and probe regressors were modeled using a canonical hemodynamic response function (HRF) with durations of 4 s (2 TR) and 9 s (4.5 TR), respectively. The cue phase modeled both full and partial trials, whereas the probe phase modeled only full trials due to the structure of the experimental design. This compound event-related approach effectively separates the HRF response for the probe phase from the cue phase (Ollinger et al. 2001). An additional regressor was also included at the onset of the first rating with a duration of 9 s (4.5 TR; total duration of three ratings), which was not of interest in the current study.

We created contrasts comparing each individual visual perspective condition to the control task (i.e., Own Eyes > Retrieve; Observer > Retrieve). We performed a conjunction analysis to identify the regions commonly activated due to retrieval orientation in the cue phase for Own Eyes and Observer, compared to the Retrieve condition (i.e., [Own Eyes > Retrieve] ∩ [Observer > Retrieve]). To assess differences between the perspective conditions, we conducted a paired sample t-test to compare Own Eyes > Retrieve and Observer > Retrieve.

This yielded two interaction contrasts:

i. (Own Eyes > Retrieve) > (Observer > Retrieve)
ii. (Observer > Retrieve) > (Own Eyes > Retrieve).

For ease of reference in the results, we will refer to these as:

i. Own Eyes > Observer & Retrieve
ii. Observer > Own Eyes & Retrieve, respectively.

The same contrast structure and analysis procedures were applied to both the cue and probe phases. Cluster-level family-wise error (FWE) correction was applied in SPM12 using Gaussian random field theory, with a cluster-defining threshold of p < .001 (uncorrected) and a cluster-level FWE threshold of p < .05. This approach, in combination with spatial smoothing, has been shown to maintain acceptable false-positive rates (Eklund et al. 2016).

#### Region of Interest (ROI) Analysis

We conducted a series of ROI analyses to test hypotheses about the role of the posterior parietal cortex and hippocampus in retrieval orientation. Anatomical ROIs for the posterior parietal cortices were defined using the JuBrain Anatomy Toolbox (Eickhoff et al. 2005). Specifically, we extracted probabilistic maps for the PGa and PGp subregions within the AG and the 7A, 7M, and 7P subregions of the precuneus, separately for left and right hemispheres. Hippocampal ROIs were defined based on probabilistic anatomical maps from NeuroVault (Collection 3731; https://identifiers.org/neurovault.collection:3731), following procedures consistent with prior research (Ritchey et al. 2015). The anterior hippocampus was defined using the head subregion, while the posterior hippocampus was defined by averaging across the body and tail subregions. These ROIs were used to examine functional differences along the anterior–posterior axis of the hippocampus. Signal intensity values were extracted based on percent signal change using MarsBar (Brett et al. 2002). ROI results are therefore independent of the contrasts used to define activation in the voxelwise group maps.

R Statistical Software (R Core Team 2025) was used for ROI analyses. To assess how retrieval orientation modulated activity within these regions, we conducted separate repeated measures ANOVAs for the cue and probe phases, separately within each hemisphere. Specifically, for the AG we ran a 2 (Region: PGa, PGp) x 3 (Retrieval Orientation: Own Eyes, Observer, Retrieve) x 2 (Hemisphere: Left, Right) repeated measures ANOVA. Similarly, for the precuneus, we conducted a 3 (Region: 7A, 7M, 7P) x 3 (Retrieval Orientation: Own Eyes, Observer, Retrieve) repeated measures ANOVA separately in left and right hemispheres. To control familywise error across the four omnibus tests, a Bonferroni-corrected significance threshold of .0125 was used. For significant omnibus effects, Holm-Bonferroni corrections were applied to all pairwise comparisons, controlling the familywise error rate across retrieval orientation conditions within each region. Repeated measures ANOVAs were corrected for violations of sphericity using Greenhouse Geisser adjustments where appropriate.

#### Brain-Behavior Relationship

One goal of the present study was to examine how brain activity during the cue and probe phases predicted AM characteristics during retrieval, as well as participants’ ability to maintain a retrieval perspective. To this end, we employed linear mixed-effects models to predict behavioral ratings during scanning from percent signal change estimated using trial-level activity derived from full-trials during the cue and probe phases of each retrieval trial (Chen et al. 2013). This approached allowed us to account for the hierarchical structure of the data, with individual memories (trials) nested within participants.

Separate linear mixed-effects models were conducted for each behavioral outcome: vividness, emotional intensity, and perspective maintenance ratings during Session 2. Each model included the following fixed effects: 1) Retrieval orientation condition (Own Eyes, Observer, Retrieve; entered as a categorical predictor), 2) Percent signal change in a given region of interest (ROI), and 3) Session 1 Initial characteristics as covariates: including vividness, emotional intensity, and a perspective bias score (own eyes minus observer ratings). Participant ID was included as a random intercept to account for repeated measures. Models were run separately for each phase (cue and probe) and for each ROI set: AG (PGa and PGp) and precuneus (7A, 7M, and 7P). Importantly, all ROIs within a given set were entered simultaneously in the same model to estimate the unique contribution of each subregion to behavioral outcomes, controlling for variance explained by the other ROIs. This approach allows us to determine which subregions exhibit distinct brain-behavior relationships, beyond shared variance across anatomically adjacent areas. As such, significant effects can be interpreted as region-specific, rather than global or overlapping across the ROI set. This approach is well-suited for testing hypotheses about distributed but functionally specialized contributions within anatomically defined networks. In contrast, treating region as a categorical factor would allow for direct statistical comparison of effect magnitudes across regions, but does not isolate each region’s unique explanatory power. Given our theoretical framework, where angular gyrus and precuneus subregions are hypothesized to support distinct, parallel functions (e.g., goal setting, visuospatial fidelity, affective regulation), our analysis was designed to test how each subregion independently contributes to the construction of subjective memory experience.

To determine whether brain-behavior relationships differed by retrieval orientation, we first tested for ROI × Condition interactions. When significant interactions were observed, we conducted simple slopes analyses to examine brain-behavior associations within each condition. In the absence of a significant interaction, we interpreted main effects of neural activity across conditions. For clarity, only results involving ROI predictors and their interactions with retrieval condition are reported in the main text. As noted by Luke (2017), correction for multiple comparisons is not required when model terms reflect distinct theoretical hypotheses rather than repeated tests of the same hypothesis. Our linear mixed-effects models were structured accordingly: we estimated separate models for each behavioral outcome (vividness, emotional intensity, and perspective maintenance), using predefined ROIs selected based on our theoretical framework. Each model tested a focused set of brain-behavior relationships, and results are reported transparently with full estimates, 95% confidence intervals, and p-values provided in Supplemental Tables 1-12.

To further assess regional specificity, we conducted exploratory linear mixed-effects models in which ROI was treated as a categorical factor (PGa vs PGp; 7A vs 7M vs 7P). These models tested Region × fMRI signal × Condition interactions allowing us to evaluate whether the strength of the brain-behavior relationship differed significantly across subregions. This approach tests a distinct hypothesis: whether a given effect is stronger in one region than another, rather than whether a region uniquely contributes to behavior after accounting for shared variance. While these models offer a complementary perspective, they do not isolate region-specific variance and are more sensitive to collinearity. Results are summarized in the Supplemental Results and Supplemental Tables 13-24.

## Results

### Behavioral Results

We first ran separate repeated measures ANOVAs for each rating from Session 1 to investigate whether there was a significant difference in the initial AM ratings across conditions. As expected, there was no significant difference in Session 1 ratings across conditions (see Supplemental Table 25 for means and SDs).

Turning to Session 2 ratings, we conducted separate repeated measures ANOVAs to investigate whether there was a difference in vividness, emotional intensity, and perspective maintenance during fMRI scanning as a function of retrieval orientation (for means and SDs, see Supplemental Table 25). The results did not reveal a significant difference in vividness, *F* (2, 58) = 2.97, *p* = .059, *η_p_^2^* = .093, or emotional intensity, *F* (2, 58) = 1.86, *p* = .164, *η_p_^2^* = .060. However, there was a significant difference in perspective maintenance as a function of retrieval orientation, *F* (1.32, 38.19) = 6.74, *p* = .008, *η_p_^2^* = .189, such that participants were better able to maintain their perspective in the Retrieve condition (i.e., their natural perspective) compared to the Observer condition (*p* = .003; see Supplemental Figure 2). There were no significant differences in reaction time (RT) of Session 2 ratings across the conditions (all *p*-values > .110).

### fMRI Results: Whole-brain Analyses

#### Cue Phase

In line with the primary goal of the present study, we aimed to isolate the neural regions contributing to retrieval orientation when participants were cued to retrieve AMs from a specific perspective or not. The results of the conjunction analysis revealed significant overlap in neural recruitment of the left AG when cued by Own Eyes and Observer perspectives compared to the Retrieve condition (see Supplemental Table 26 and Figure 2A). This finding aligns with previous research highlighting the role of AG when adopting a particular visual perspective during AM retrieval (St Jacques et al. 2017; St. Jacques et al. 2018), but extends this research by demonstrating that the AG is driven by prompts to adopt a specific perspective rather than the products of memory retrieval. The pattern of results in the AG remained unchanged after controlling for individual differences perspective bias score (own eyes minus observer ratings), although visual cortex effects were reduced.

**Figure 2.**
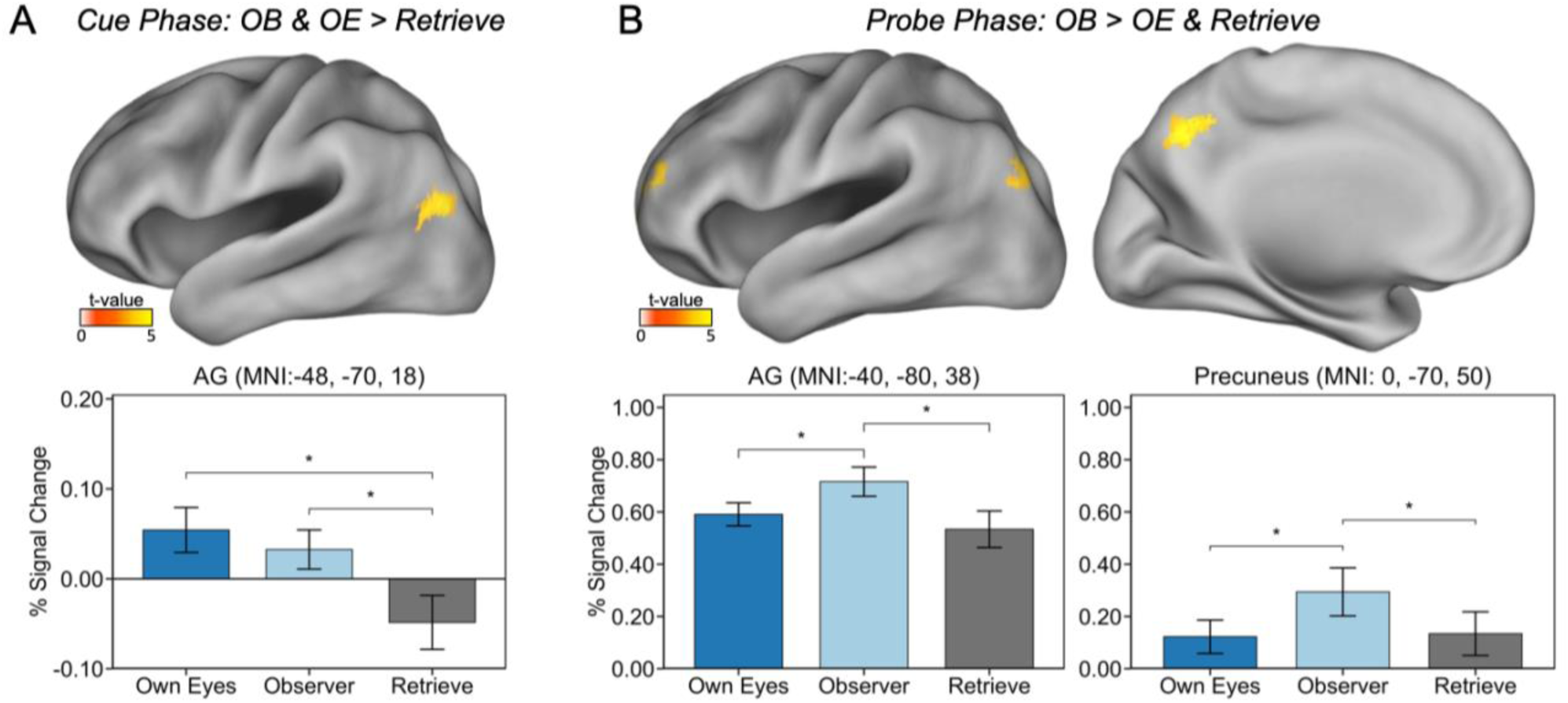
Whole-brain analysis. (A) During the cue phase, a conjunction analysis revealed common neural recruitment in the angular gyrus (AG) for Own Eyes and Observer cues relative to Retrieve cues. (B) During the probe phase, the Observer condition elicited greater activation in both the AG and precuneus compared to the Own Eyes and Retrieve conditions. Graphs represent the percent signal change extracted based on spherical ROIs centered around the peak of the response from the whole-brain analysis MarsBar (Brett et al. 2002). Asterisks in the figure reflect the direction of whole-brain contrasts and do not represent independent statistical tests.

Turning to differences in the cue phase based on visual perspective, we found greater recruitment in bilateral visual cortex (BA 18) in the Observer, compared to the Own Eyes and Retrieve cues (see Supplemental Table 27). However, there were no regions with greater recruitment in Own Eyes versus Observer and Retrieve cues.

#### Probe Phase

The conjunction analyses indicated that there were no overlapping regions in the Own Eyes and Observer versus Retrieve condition. There were also no differences in Own Eyes compared to the Observer and Retrieve condition. Importantly, the results revealed greater neural recruitment in the Observer compared to Own Eyes and Retrieve condition in both the left AG and precuneus (see Figure 2B). There was also greater recruitment in bilateral anterior prefrontal cortex (see Supplemental Table 27). Thus, during the probe phase, adopting an observer-like perspective during AM retrieval continued to require the left AG, along with additional recruitment of the precuneus and anterior prefrontal cortex. The pattern of results remained unchanged, after controlling for individual differences perspective bias score (own eyes minus observer ratings).

### ROI Analyses: Posterior Parietal Cortices

We conducted targeted ROI analyses examining the contribution of posterior parietal subregions of the AG (PGa and PGp) and precuneus (7A, 7M, 7P) for cue and probe phases, separately, using anatomically defined regions based on cytoarchitectonic probability maps (Eickhoff et al., 2005). Given prior research suggesting hemispheric differences in the AG (for review see St. Jacques, 2015) we included left and right hemisphere as a factor when analyzing AG subregions (PGa and PGp). In contrast, we examined bilateral effects in the precuneus by averaging across hemispheres in precuneus ROIs (7A, 7M, 7P).

#### Cue Phase

In AG, a significant Hemisphere x Retrieval Orientation x Region interaction (F(2, 58) = 8.837, p < .001, ηp² = .23) during the cue phase indicated that preparatory retrieval orientation effects differed across AG subregions as a function of hemisphere. Follow-up analyses conducted separately within each hemisphere revealed a Retrieval Orientation x Region interaction in the left AG (F(2, 58) = 12.708, p < .001, ηp² = .30). In the left PGa, there was no effect of retrieval orientation (all p’s > .381). In contrast, left PGp activation was greater for both Own Eyes and Observer (p = .019 and p = .007, respectively) compared to Retrieve cues, with no difference between Own Eyes and Observer. Although there was also a significant Retrieval Orientation x Region interaction in the right AG (F(2, 58) = 6.191, p = .004, ηp² = .18), simple effects were not significant after Holm-Bonferroni correction (see Supplemental Figure 3). Together, these findings indicate that preparatory perspective cues selectively engage left PGp, supporting retrieval orientation prior to memory access.

**Figure 3.**
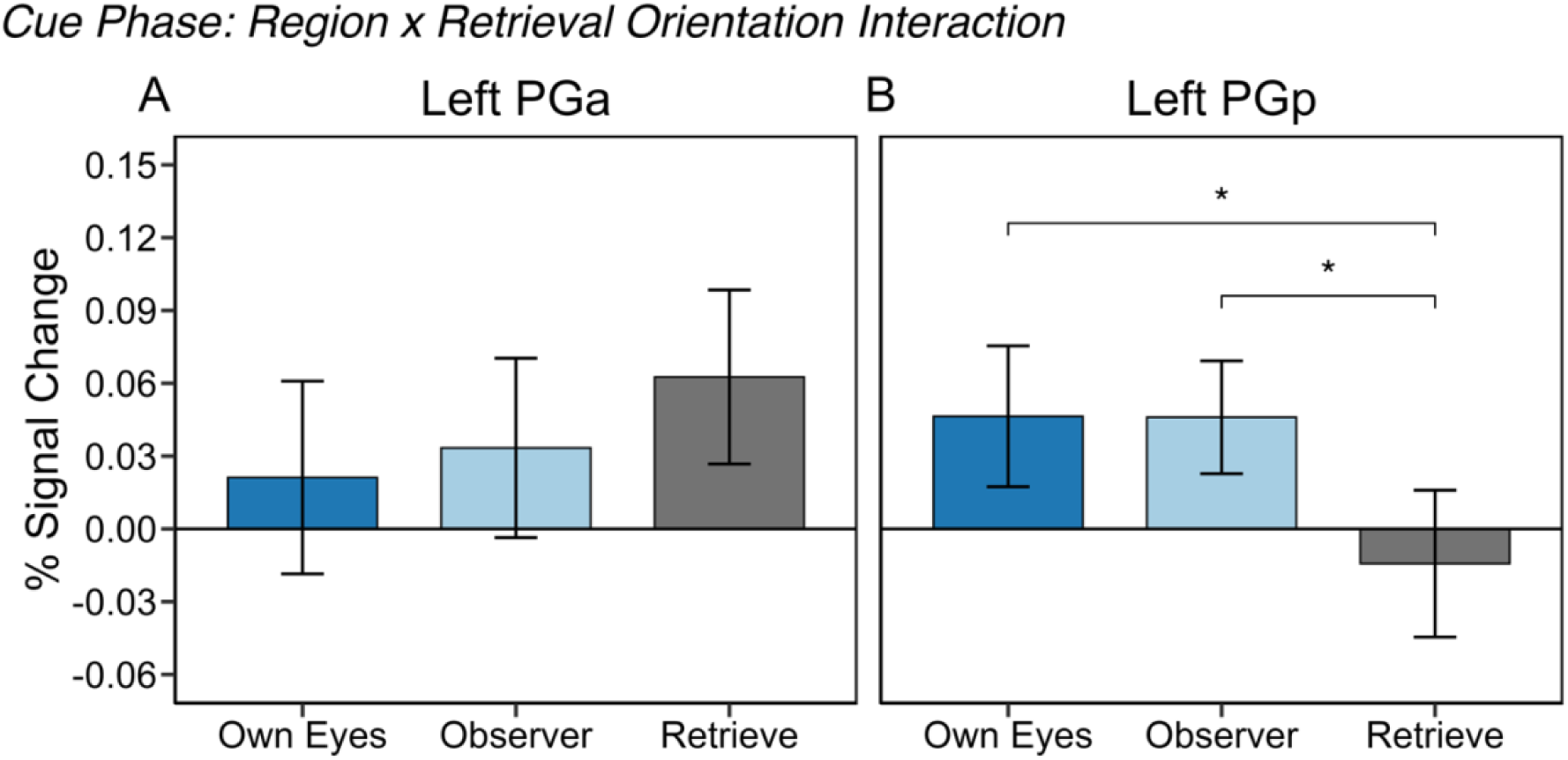
ROI analysis in left angular gyrus (AG). During the cue phase, a significant Region x Retrieval Orientation interaction indicated no significant differences in retrieval orientation in left PGa (A), but in left PGp (B) there was stronger activation for Own Eyes and Observer compared to Retrieve cues.

In the precuneus (collapsed across hemisphere), there was a significant main effect of Region (F(2, 58) = 12.708, p < .001, ηp² = .69) reflecting overall differences in activation across precuneus subregions. Holm-corrected pairwise comparisons collapsed across retrieval orientation indicated that 7M (M = .379, SD = .286) exhibited greater activation than both 7A (M = −.118, SD = .176; p < .001) and 7P (M = .024, SD = .217; p < .001), and that 7P was also more active than 7A (p = .0004). Thus, precuneus activation followed a graded pattern (7M > 7P > 7A) that was consistent across retrieval orientation conditions. Although there was a main effect of Retrieval Orientation (F(1.88, 54.42) = 4.60, p = .016, ηp² = .14), this effect did not survive the omnibus corrected threshold (α = .0125) and was therefore not interpreted further. The Retrieval Orientation × Region interaction was also not significant (F(2.82, 81.73) = 2.04, p = .118, ηp² = .07) indicating no differences in retrieval orientation across precuneus regions.

#### Probe Phase

In AG, there were main effects of Hemisphere (F(1, 29) = 97.03, p < .001, ηp² = .77) and Region (F(1, 29) = 31.15, p < .001, ηp² = .52), as well as a significant Hemisphere x Region interaction (F(2, 58) = 8.982, p = .006, ηp² = .24). Follow-up analyses decomposing the Hemisphere × Region interaction revealed greater activation in PGp than PGa in both hemispheres. This region effect was robust in the right hemisphere (p < .001), whereas the corresponding effect in the left hemisphere was weaker (p = .019). The main effect of Retrieval Orientation did not meet the omnibus corrected threshold, F(2, 58) = 5.57, p = .015, ηp² = .16, nor did interactions with Retrieval Orientation (all ps > .035). Accordingly, retrieval orientation effects were not robust at the ROI level following correction for multiple comparisons.

In the precuneus, there was a significant main effect of Region (F(1.34, 38.81) = 38.18, p < .001, ηp² = .57) indicating substantial differences in overall activation across precuneus subregions during the probe phase. Holm-corrected pairwise comparisons collapsed across retrieval orientation indicated that 7M (M = .535, SD = .541) exhibited greater activation than both 7A (M = −.171, SD = .312; p < .001) and 7P (M = .103, SD = .403; p < .001), and that 7P was also more active than 7A (p < .001). Thus, similar to the cue phase, precuneus activation followed a graded pattern (7M > 7P > 7A) that was consistent across retrieval orientation conditions during the probe phase. The main effect of Retrieval Orientation did not meet the omnibus corrected threshold (F(1.96, 56.74) = 3.24, p = .047, ηp² = .10) nor did the Retrieval Orientation × Region interaction (F(2.83, 81.95) = 2.71, p = .054, ηp² = .09). Accordingly, retrieval orientation effects in the precuneus were not robust after correction for multiple comparisons.

#### Summary

Taken together, the ROI analyses revealed a selective effect of retrieval orientation during the cue phase in the AG, localized to left PGp, where both Own Eyes and Observer cues elicited greater activation than Retrieve cues. No corresponding retrieval orientation effects were observed in PGa or in the precuneus, which instead showed a stable subregional hierarchy (7M > 7P > 7A) that was invariant across retrieval orientation conditions. Importantly, whole-brain analyses revealed greater recruitment of AG and precuneus during observer-perspective retrieval, suggesting that retrieval orientation effects during the probe phase may be detectable at the regional level but do not map cleanly onto subregional dissociations under stringent correction. Whole-brain clusters corresponding to these effects overlapped primarily with PGp in the AG (57.6% of voxels) and modestly with 7A (4.9%) and 7P (3.2%) in the precuneus. Relative to the active baseline, 7A was consistently deactivated across phases, whereas 7P was near baseline during the cue but showed increased activation during the probe phase.

### Brain-Behavior Relationship

To assess how neural recruitment during the cue and probe phases relates to AM characteristics, we examined associations between percent signal change in the posterior parietal ROIs and ratings of vividness, emotional intensity, and perspective maintenance acquired on each fMRI retrieval trial. Based on our whole-brain results, which revealed significant activation in the left angular gyrus but midline precuneus, and supported by the targeted ROI analyses, we focused on left-lateralized AG subregions (PGa and PGp) and averaged across hemispheres for precuneus ROIs (7A, 7M, 7P). This approach reflects the functional lateralization observed for AG, particularly the selective sensitivity of left PGp to perspective cues during both cue and probe phases, and the more bilaterally distributed or midline engagement of the precuneus across retrieval conditions. As mentioned earlier, we reported only results involving interactions between neural activity in the hypothesized ROIs and retrieval orientation conditions for clarity. If there was no significant interaction, the ROI main effects were interpreted.

#### Vividness

We first tested whether vividness ratings were predicted by neural activation in the AG during the cue and probe phase. As expected, during the cue phase none of the subregions in the left AG predicted vividness ratings (left PGa: β = −.015, SE = .039, 95% CI [−.09, .06], t (2066) = −.40, p = .696; left PGp: β = −.065, SE = .046, 95% CI [−.16, .03], t (2060.1) = −1.41, p = .159). Although there was a significant left PGa and retrieval orientation interaction (β = .182, *SE* = .071, 95% CI [.04, .32], *t* (2063.7) = 2.56, *p* = .010), simple slope estimations did not reveal a significant relationship between left PGa activation during the cue phase and vividness ratings. During the probe phase, neural recruitment in left PGa activation did not significantly predict vividness ratings (*β* = −.014, *SE* = .043, 95% CI [−.10, .07], *t* (2047.9) = −.33, *p* = .744). However, neural recruitment in the left PGp activity during the probe phase was a significant predictor of vividness ratings (*β* = .109, *SE* = .049, 95% CI [.01, .21], *t* (1933) = 2.22, *p* = .027), reflecting that left PGp contributes to the vivid reconstruction of memory content (see Figure 5A). No significant interaction was observed between left PGp and retrieval orientation conditions during the probe phase.

**Figure 5.**
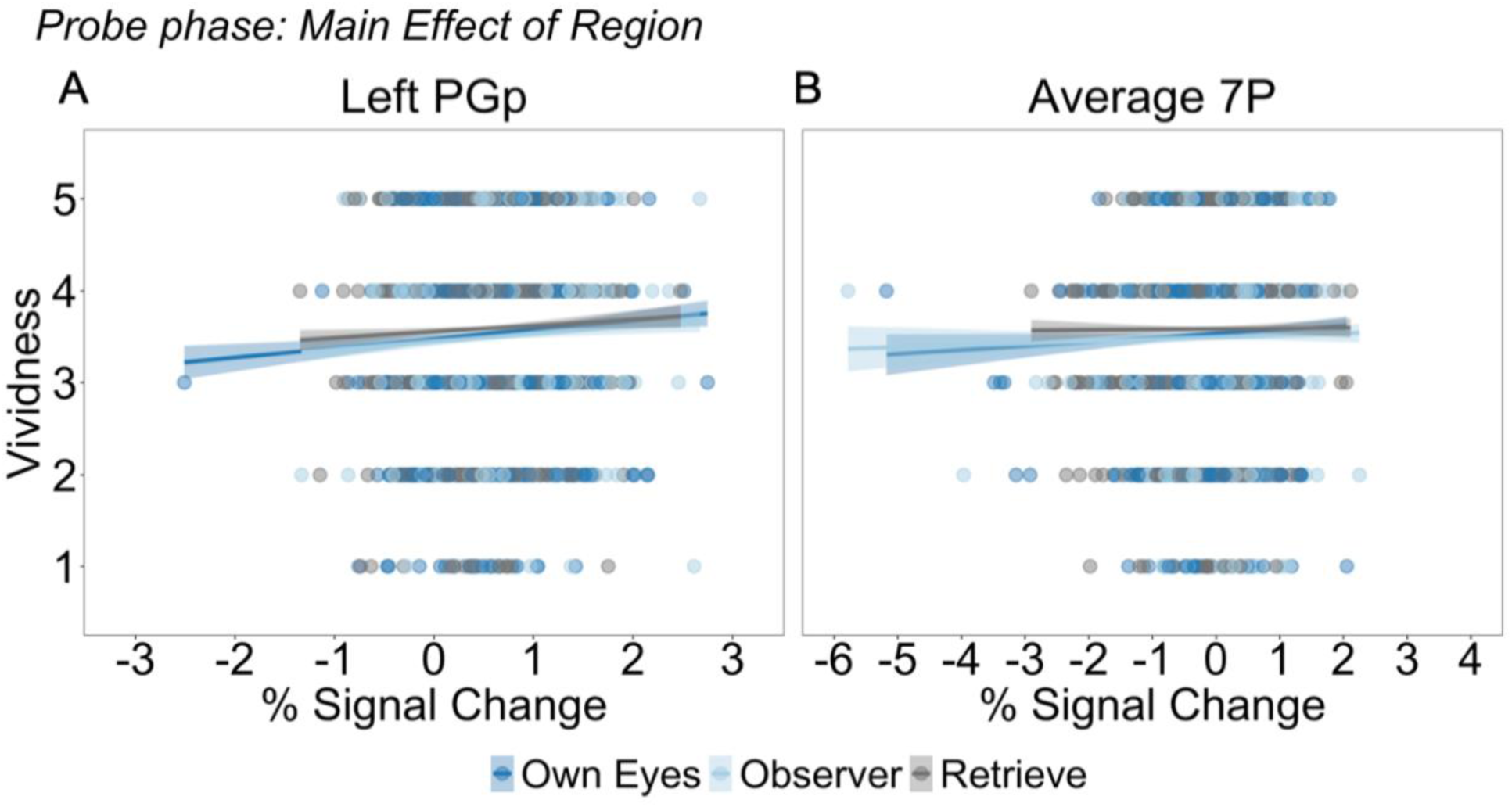
Predicted vividness ratings (1 = low, 5 = high) as a function of % signal change during the probe phase. Interactions were tested separately for AG (PGa, PGp) and precuneus (7A, 7M, 7P). Lines represent estimated values reflecting a Main Effect of Region in Left PGp (A) and Average 7P (B). In both regions, there was a stronger positive association between signal change and vividness. *S*haded areas indicating 95% confidence intervals

Next, we examined whether brain activity in precuneus subregions during the cue and probe phases predicted vividness ratings. During the cue phase, none of the precuneus subregions significantly predicted vividness ratings (7A: *β* = −.041, *SE* = .058, 95% *CI* [−.16, .07], *t* (2072.2) = −.71, *p* = .477; 7M: *β* = −.023, *SE* = .034, 95% CI [−.09, .04], *t* (2071.9) = −.67, *p* = .502); 7P: *β* = .035, *SE* = .049, 95% CI [−.06, .13], *t* (2069.7) = .70, *p* = .482). There were also no significant interactions. During the probe phase, neither 7A or 7M recruitment significantly predicted vividness ratings (7A: *β* = −.029, *SE* = .063, 95% CI [−.15, .09], *t* (2041.8) = −.46, *p* = .643; 7M: *β* = −.038, *SE* = .032, 95% CI [−.10, .03], *t* (2046.3) = −1.17, *p* = .243). However, there was a significant positive relationship between 7P activity and vividness ratings during the probe phase (*β* = .132, *SE* = .052, 95% CI [.03, .23], *t* (2070.2) = 2.53, *p* = .011), reflecting that stronger engagement of 7P contributed to more vivid remembering (see Figure 5B). There was no significant interaction between 7P activity and retrieval orientation.

#### Emotional Intensity

During the cue phase, none of the subregions in the AG significantly predicted emotional intensity (left PGa: *β* = −.063, *SE* = .040, 95% CI [−.14, .02], *t* (2024.5) = −1.54, *p* = .124; left PGp: *β* = .057, *SE* = .047, 95% CI [−.04, .15], *t* (2022) = 1.22, *p* = .224). Similarly, during the probe phase, AG subregions were not significant predictors of emotional intensity (left PGa: *β* = .077, *SE* = .044, 95% CI [−.01, .17], *t* (2031.3) = 1.74, *p* = .081; left PGp: *β* = .027, *SE* = .050, 95% CI [−.07, .13], *t* (1956.4) = .54, *p* = .591). There were no significant interactions between left AG subregions and retrieval orientation in the cue and probe phases.

Next, we examined whether brain activity in precuneus subregions predicted emotional intensity ratings. None of the subregions in the precuneus during the cue phase were significant predictors of emotional intensity (7A: *β* = −.040, *SE* = .060, 95% *CI* [−.16, .08], *t* (2008.4) = −.66, *p* = .509; 7M: *β* = −.042, *SE* = .034, 95% *CI* [−.11, .03], *t* (2029.9) = −1.22, *p* = .221; 7P: *β* = .048, *SE* = .051, 95% *CI* [−.05, .15], *t* (2037.7) = .95, *p* = .342). There was no significant interaction between 7A and retrieval orientation. During the probe phase, neither 7A (*β* = −.038, *SE* = .063, 95% CI [− .16, .09], *t* (1995.6) = −.61, *p* = .544) or 7P (*β* = .069, *SE* = .053, 95% CI [−.04, .17], *t* (2011.3) = 1.30, *p* = .194) were significant predictors of emotional intensity. However, 7M activation during the probe phase was negatively associated with emotional intensity ratings (*β* = −.084, *SE* = .033, 95% CI [−.15, −.02], *t* (1970.3) = −2.53, *p* = .012), reflecting that greater 7M recruitment predicted less emotional intensity during AM retrieval. Importantly, there was also a significant interaction between retrieval orientation and 7M activation during the probe phase, specifically when comparing the Observer and Retrieve conditions (*β* = −.118, *SE* = .053, 95% CI [−.22, −.01], *t* (1985.1) = −2.23, *p* = .026). To follow up the significant Retrieval Orientation × 7M activation interaction during the probe phase, we estimated simple slopes of 7M activation on emotional intensity within each retrieval orientation condition. A significant negative slope was observed in the Observer condition (*β* = −.153, *SE* = .045, 95% *CI* [−.24, −.06], *t* (2017) = −3.35, *p* < .001) indicating that greater 7M recruitment was associated with reduced emotional intensity. The corresponding slopes in the Retrieve (*β* = −.035, *SE* = .045, 95% *CI* [−.12, .05], *t* (2025) = −.78, *p* = .437) and Own Eyes (*β* = −.063, *SE* = .044, 95% *CI* [−.15, .02], *t* (2016) = −1.43, *p* = .154) conditions were not statistically significant. In contrast, there were no interactions between 7M activity when comparing the Observer versus Own Eyes conditions (*β* = −.090, *SE* = .052, 95% *CI* [−.19, .91], *t* (1988.9) = −1.74, *p* = .083) or the Own Eyes versus Retrieve conditions (*β* = −.028, *SE* = .051, 95% *CI* [−.13, .07], *t* (1999) = −.54, *p* = .587). This pattern indicates that adopting an observer perspective uniquely engages 7M processes that dampen emotional intensity (see Figure 6).

**Figure 6.**
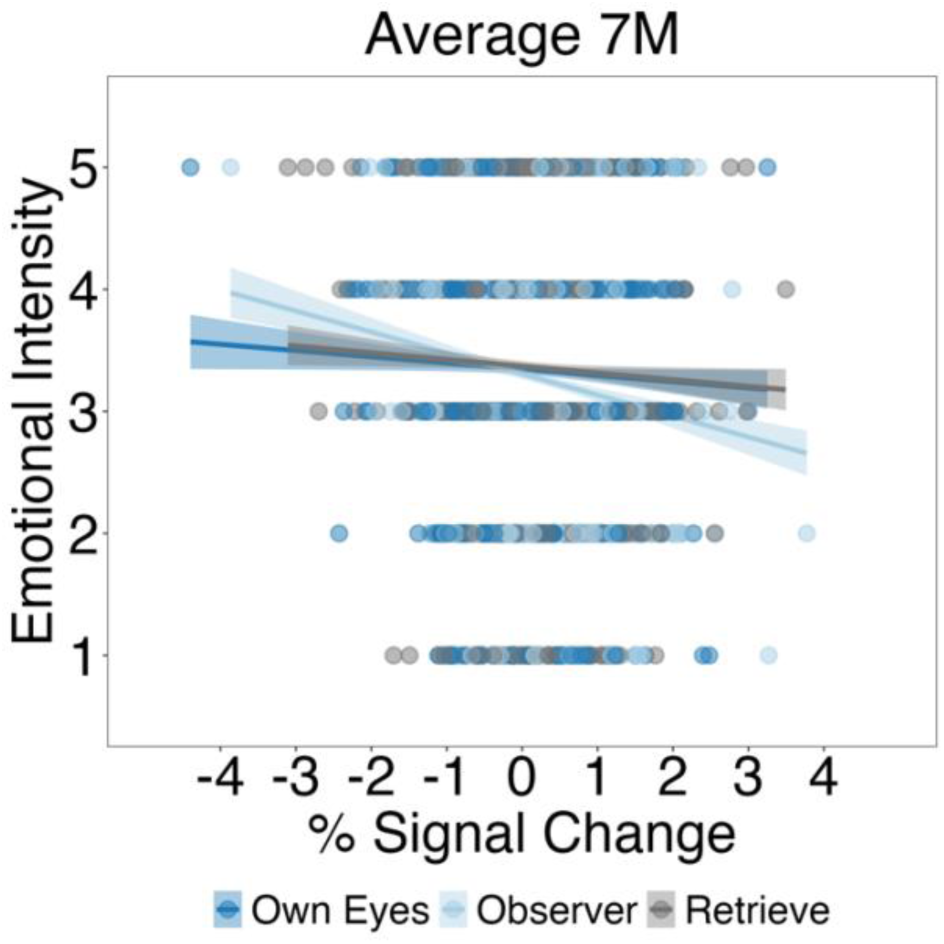
Predicted emotional intensity ratings (1 = low, 5 = high) as a function of % signal change during the probe phase. Lines represent estimated values from Region × Retrieval Orientation interaction in 7M. There was a stronger negative association between signal change and emotional intensity in the Observer condition compared to the Retrieve condition. Shaded areas indicating 95% confidence intervals.

#### Perspective Maintenance

During the cue phase, neural recruitment in left PGa significantly predicted perspective maintenance ratings (*β* = .098, *SE* = .050, 95% CI [.00, .19], *t* (2049.5) = 1.97, *p* = .049). Critically, there was a significant interaction between left PGa and retrieval orientation, when comparing the Own Eyes and Observer conditions (*β* = −.209, *SE* = .090, 95% CI [−.39, −.03], *t* (2035.8) = −2.31, *p* = .021) and Own Eyes and Retrieve conditions (*β* = −.238, *SE* = .090, 95% CI [−.42, −.06], *t* (2041.4) = −2.63, *p* = .009), but not when comparing the Observer and Retrieve conditions (*β* = −.029, *SE* = .088, 95% CI [−.20, .14], *t* (2040) = −.33, *p* = .741). To follow up the significant Retrieval Orientation × left PGa activation interaction during the cue phase, we estimated simple slopes of left PGa activation on perspective maintenance ratings within each retrieval orientation condition. There was a significant positive slope in the Own Eyes condition (*β* = .247, *SE* = .074, 95% CI [.10, .39], *t* (2052) = 3.33, *p* < .001), reflecting that higher activity in left PGa during the cue phase contributed to the ability to maintain an own eyes perspective (see Figure 7A). However, the slopes were not significant in either the Observer condition (*β* = .038, *SE* = .071, 95% CI [−.10, .18], *t* (2053) = .55, *p* = .584) or Retrieve condition (*β* = .009, *SE* = .071, 95% CI [−.13, .15], *t* (2045) = .14, *p* = .892). In contrast, during the cue phase left PGp was not a significant predictor of perspective maintenance ratings (*β* = −.088, *SE* = .058, 95% CI [−.20, .03], *t* (2051.6) = −1.51, *p* = .131). During the probe phase, left PGa did not significantly predict perspective maintenance (*β* = −.095, *SE* = .055, 95% CI [−.20, .01], *t* (2070.4) = −1.72, *p* = .086). However, left PGp activation during the probe phase was positively associated with perspective maintenance, *β* = .145, *SE* = .062, 95% CI [.02, .27], *t* (1998.2) = 2.33, *p* = .020, reflecting that higher left PGp activation contributed to more stable perspectives during AM retrieval. There were no significant interactions between PGp and retrieval orientation in the probe phase.

**Figure 7.**
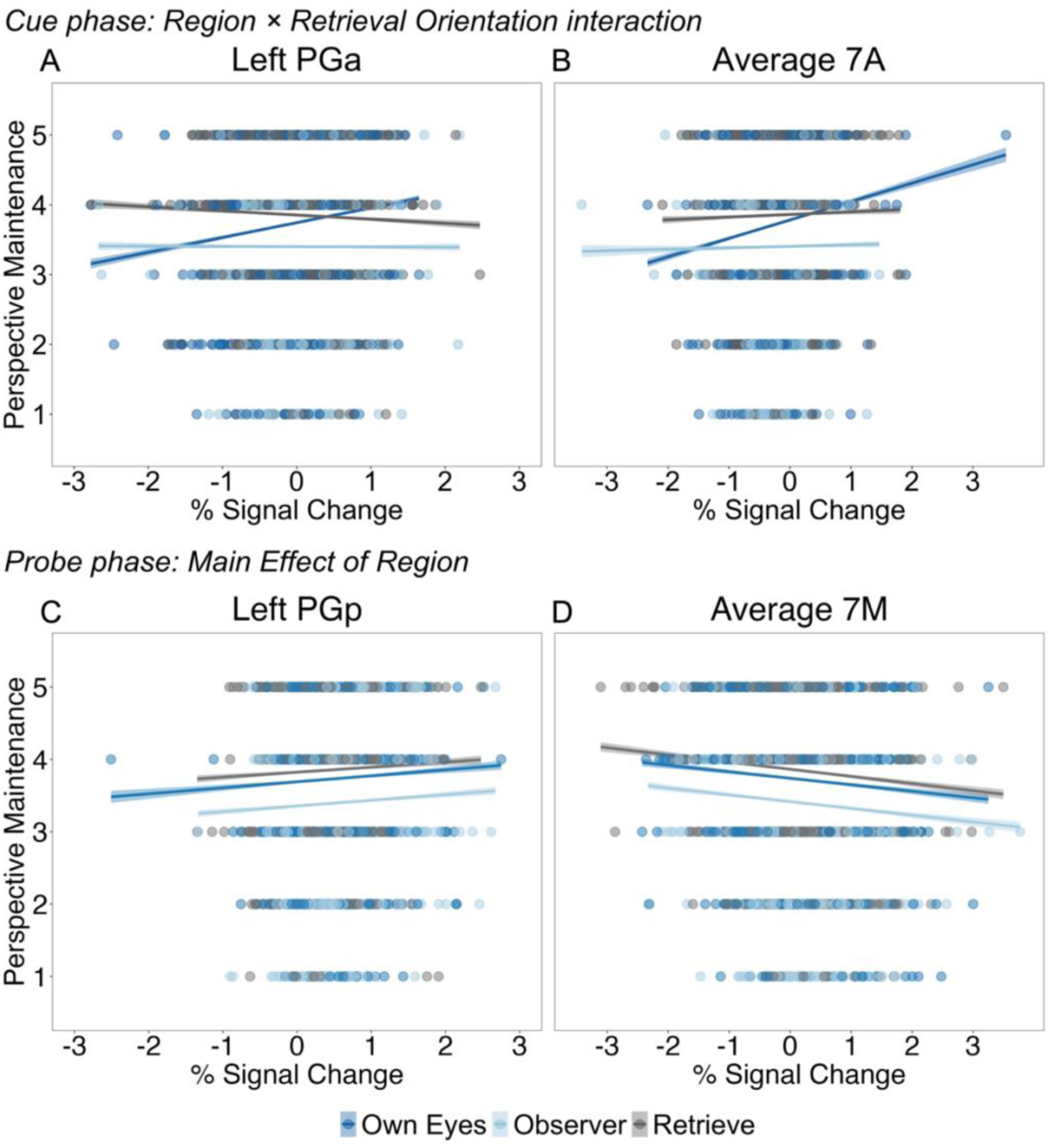
Predicted perspective maintenance ratings (1 = low, 5 = high) as a function of % signal change during the cue and probe phase. Interactions were tested separately AG (PGa, PGp) and precuneus (7A, 7M, 7P). Lines represent estimated values. During the cue phase, there was a significant Region × Retrieval Orientation interactions in Left PGa (A) and Average 7A (B), reflecting a stronger positive association between signal change and perspective maintenance in the Own Eyes condition compared to the Observer and Retrieve conditions. During the probe phase, there was a significant Main Effect of Region in Left PGp (C), reflecting a positive association between signal change and perspective maintenance. In contrast, in Average 7M (D) there was a negative association between signal change and perspective maintenance. Shaded areas indicating 95% confidence intervals. One visually identified extreme value in the Own Eyes condition for 7A did not influence the results and was retained in both the analysis and the figure.

Turning to the precuneus, during the cue phase there was no significant relationship between 7M (*β* = −.020, *SE* = .043, 95% CI [−.11, .06], *t* (2067.9) = −.49, *p* = .627), and 7P recruitment, (*β* = −.058, *SE* = .062, 95% CI [−.18, .06], *t* (2067) = −.93, *p* = .352). However, 7A activation in the cue phase was significantly related to perspective maintenance, *β* = .167, *SE* = .073, 95% CI [.02, .31], *t* (2060.1) = 2.28, *p* = .023. Importantly, this effect was qualified by a significant 7A and condition interaction, specifically when comparing the Own Eyes condition to the Observer condition, *β* = −.224, *SE* = .104 95% CI [−.43, −.02], *t* (2036.8) = −2.14, *p* = .032, and Retrieve conditions, *β* = −.214, *SE* = .103, 95% CI [−.42, −.01], *t* (2029.2) = −2.06, *p* = .039. However, Observer and Retrieve conditions were not significantly different from each other, *β* = .009, *SE* = .106, 95% CI [−.20, .22], *t* (2037.1) = .09, *p* = .926. To follow up the significant Retrieval Orientation × 7A activation interaction during the cue phase, we estimated simple slopes of 7A activation on perspective maintenance ratings within each retrieval orientation condition. There was a significant positive slope in the Own Eyes condition, *β* = .313, *SE* = .093, 95% CI [.13, .50], *t* (2055) = 3.37, *p* < .001, indicating that higher activity in the 7A during the cue phase was associated with a better ability to maintain an own eyes perspective (see Figure 7B). There were no significant slopes in either the Observer condition, *β* = .089, *SE* = .096, 95% CI [−.10, .28], *t* (2059) = .93, *p* = .355, or Retrieve condition, *β* = .099, *SE* = .096, 95% CI [−.09, .29], *t* (2054) = 1.04, *p* = .301.

During the probe phase, we observed a significant negative association between 7M recruitment and perspective maintenance (*β* = –.183, *SE* = .040, 95% *CI* [–.26, –.10], *t*(2028.2) = –4.48, *p* < .001), reflecting that greater 7M activity tracked lower perspective maintenance ratings. In contrast, 7P recruitment was positively associated with perspective maintenance (*β* = .151, *SE* = .065, 95% *CI* [.02, .28], *t*(2052.6) = 2.30, *p* = .022). Although 7A recruitment was not significantly related to perspective maintenance when collapsed across conditions (*β* = .061, *SE* = .079, 95% *CI* [–.09, .22], *t*(2035.9) = 0.77, *p* = .442), a significant interaction emerged between 7A activity and retrieval orientation condition. Follow-up analyses revealed a significant positive association between 7A recruitment and perspective maintenance in the Own Eyes condition (*β* = .308, SE = .122, 95% CI [.07, .55], *t*(2068) = 2.54, p = .011), indicating that higher 7A activation during memory retrieval supports the ability to maintain an own-eyes perspective. Crucially, this effect was significantly stronger than in the Observer (*β* = –.410, *SE* = .167, 95% *CI* [–.74, –.08], *t*(2029.4) = –2.45, *p* = .015) and Retrieve (*β* = –.333, *SE* = .161, 95% *CI* [–.65, –.02], *t*(2030.3) = –2.07, *p* = .039) conditions. In both of those conditions, the relationship between 7A recruitment and perspective maintenance was non-significant (Observer: *β* = –.101, *SE* = .130, 95% *CI* [–.36, .15], *t*(2064) = –0.78, *p* = .436; Retrieve: *β* = –.024, *SE* = .121, 95% *CI* [–.26, .21], *t*(2066) = –0.202, *p* = .840). The Observer vs. Retrieve contrast was also not significant (*β* = .076, SE = .167, 95% *CI* [–.25, .41], *t*(2027.2) = 0.46, *p* = .647). No significant interaction observed between 7P activity and retrieval orientation.

#### Summary of Brain-Behavior Relationships

These results reveal temporally and functionally distinct contributions of posterior parietal subregions to the subjective characteristics of AM. During the cue phase, perspective maintenance for own-eyes retrieval was supported by activity in left PGa and 7A, highlighting their role in orienting to a first-person perspective prior to memory access. During the probe phase, activity in left PGp and 7P was positively associated with both vividness and perspective maintenance across conditions, suggesting that these posterior regions contribute to the integration of phenomenological richness and perspective coherence during retrieval. In contrast, greater 7M activity was associated with reduced perspective maintenance across conditions and lower emotional intensity specifically during observer-perspective retrieval. This pattern may reflect increased effort or instability in maintaining a coherent viewpoint, such that individuals fluctuate between perspectives rather than sustaining the instructed one. In the observer condition, this instability may promote emotional distancing, blunting the intensity of the remembered experience. Notably, 7A continued to track perspective maintenance for own-eyes retrieval during the probe phase, suggesting a sustained role in supporting a first-person viewpoint. Taken together, these findings suggest a functional division in which AG subregions support goal setting and perspective alignment, while precuneus subregions dynamically support the reconstruction of visual, emotional, and perspectival aspects of memory.

## Discussion

This study demonstrates that visual perspective cues act as retrieval orientation signals, engaging posterior parietal regions prior to memory access. A key strength of the design is the temporal separation of cue and probe, which allowed us to isolate preparatory retrieval orientation from mnemonic content during retrieval (Ollinger et al. 2001). This extends prior work on retrieval orientation in AM, which has focused on conceptual versus perceptual goals (Gurguryan and Sheldon 2019), by examining how visual perspective shapes retrieval. Unlike prior goal-based manipulations, orienting retrieval toward own eyes versus observer viewpoints targets the constructive mode of recollection itself—modulating the spatial framework, self-location, and phenomenological experience of memory. By leveraging cytoarchitectonic ROIs, we further advance the field by examining anatomically defined subregions in the AG and precuneus, allowing us to test for functional differentiation during retrieval. When participants were cued to adopt a visual perspective prior to retrieval, the left PGp subregion of the AG showed selective engagement, reflecting preparatory rather than mnemonic processes. In contrast, during the probe phase whole-brain analysis revealed elevated activation in AG and precuneus for observer-perspective retrieval, suggesting sustained recruitment of these regions when reconstructing memories from a non-default, third-person viewpoint. Although ROI analyses did not show robust subregional differentiation during the probe phase after correction. Moreover, during the probe phase, activity in PGp and 7P was positively associated with both vividness and perspective maintenance, suggesting a role in sustaining the richness and coherence of memory retrieval. In contrast, 7M activity was negatively associated with perspective maintenance across conditions and with emotional intensity specifically during observer-perspective retrieval, potentially reflecting increased effort or instability in maintaining a detached viewpoint. These findings demonstrate that perspective-specific retrieval orientations can be neurally instantiated before memory content is accessed. This cue-probe dissociation underscores the importance of distinguishing preparatory processes from memory reconstruction, with PGp emerging as a key hub for initiating viewpoint-specific memory retrieval.

The present findings suggest that perspective cues triggered preparatory retrieval orientation processes, even before memory content was accessed. While participants had not yet received the memory title during the cue phase, prior work demonstrates that individuals can proactively configure goal states in anticipation of retrieval demands (e.g., Gilmore et al., 2018; Morcom and Rugg, 2012). Similar logic underlies many retrieval orientation studies in which participants are cued to prepare to retrieve words or pictures (e.g., Herron and Rugg, 2003) or perceptual versus conceptual information (Gurguryan and Sheldon 2019). In this context, visual perspective functioned as a top-down retrieval orientation—biasing participants to adopt a particular spatial vantage point, rather than mentally simulating the full scene in advance. Our finding that cue-phase activity in PGp and 7A predicted subsequent perspective maintenance further supports the view that these signals reflect intentional goal setting rather than stimulus-driven memory reactivation.

We initially hypothesized that PGa would be recruited during the cue phase, given its proposed role in attentional orienting and goal specification (Seghier 2013). Contrary to this prediction, the most robust cue-related activation emerged in PGp, which responded more strongly to both Own Eyes and Observer cues relative to Retrieve cues. This pattern was bilateral and suggests a broader role for PGp in setting the stage for retrieval of memory content. However, PGa was not entirely uninvolved: it positively predicted perspective maintenance, particularly for the Own Eyes condition, indicating a more specialized role in aligning retrieval goals with egocentric reference frames (Binder and Desai 2011). These findings suggest a functional division between AG subregions, with PGp supporting generalized episodic preparation and reconstruction, and PGa contributing to perspective-specific goal alignment.

The cue-phase results in the precuneus further suggest a subregional specialization that complements this AG dissociation. Activation in 7A, particularly during Own Eyes trials, was positively associated with perspective maintenance, suggesting a role in specifying retrieval goals in an egocentric space (Cavanna and Trimble 2006). By contrast, 7M showed strong cue- related engagement across all conditions, consistent with a general role in episodic memory retrieval (Margulies et al. 2009). Although these effects did not emerge as a robust interaction at the ROI level, this pattern is consistent with the idea that 7A supports goal-driven perspective alignment, whereas 7M contributes more broadly to preparatory retrieval processes. Together, the findings suggest a broader organizational principle across the posterior parietal cortex, in which anterior subregions help specify retrieval goals, and more posterior/medial areas support episodic memory processes.

During the probe phase, posterior parietal regions uniquely contributed to the adopted perspective during AM retrieval. Observer-perspective retrieval elicited increased activation across AG, precuneus, posterior cingulate, and anterior prefrontal cortex, reflecting the additional cognitive demands of transforming spatial reference frames, monitoring self-location, and regulating affect (Bergouignan et al. 2014; Iriye and St Jacques 2020). Notably, PGp activation was positively associated with both vividness and perspective maintenance, reinforcing its central role in the phenomenological construction of remembered experience. In the precuneus, 7P showed enhanced activation during observer-perspective trials and predicted both vividness and perspective stability, consistent with its role in visuospatial coordination (Cavanna and Trimble 2006). Relative to the active baseline, 7A was consistently deactivated, whereas 7P showed modest above-baseline especially during the probe phase. This pattern reinforces the view that 7P contributes to mnemonic reconstruction when visuospatial perspective demands are high. In contrast, 7M activity was negatively associated with perspective maintenance across conditions, and predicted reduced emotional intensity specifically during observer-perspective retrieval. This pattern suggests that 7M may contribute to affective distancing and viewpoint instability, potentially reflecting increased demands in managing self-location or fluctuating perspectives (St Jacques et al. 2017). Together, these findings suggest functionally distinct contributions of precuneus subregions during retrieval, with 7P supporting visuospatial aspects of memory and 7M contributing to the modulation of emotional experience.

Although the hippocampus is a core region in episodic memory, we did not observe perspective-specific modulation along its long axis. Instead, activation was dominated by a robust anterior > posterior gradient, with anterior regions more engaged during the probe phase across all retrieval conditions. This pattern aligns with previous work suggesting that the anterior hippocampus supports gist-based, integrative, and affect-laden aspects of memory, whereas the posterior hippocampus is more involved in fine-grained spatial and contextual detail (Brunec et al. 2018; Poppenk et al. 2013). The absence of differential recruitment by visual perspective may reflect the fact that our design emphasized preparatory perspective adoption via parietal regions, while hippocampal engagement was more closely tied to episodic construction itself, regardless of the adopted perspective. It is also possible that parietal regions assumed a more prominent role in coordinating spatial perspective shifts under top-down control, reducing the demand on posterior hippocampal mechanisms (Maguire and Mullally 2013). Additionally, the more remote nature of the memories in our study (i.e., > 5 years) may have attenuated the spatial specificity typically attributed to the posterior hippocampus. Prior research suggests that the hippocampus may become functionally decoupled from specific perceptual details as memories age (Barry and Maguire 2019), shifting reliance toward broader schematic representations supported by anterior regions and neocortical networks. Our findings contribute to this evolving view by showing that, during remote AM retrieval, anterior hippocampal activity dominates, likely reflecting its role in integrating affective and conceptual components of autobiographical events. Future work could explore whether recent versus remote memories, or those that require active perspective shifting, differentially engage the hippocampal axis.

The dissociable contributions of PGp, 7P, and 7M suggest a division of labor across the posterior parietal cortex during perspective-guided memory retrieval (for review see St. Jacques, 2025). Specifically, PGp and 7P supported vividness and perspective maintenance during the probe phase, while PGp was also recruited during the cue phase, indicating a broader role in retrieval preparation. PGa, in contrast, predicted own-eyes perspective maintenance during the cue phase, highlighting its contribution to spatial perspective setting. Notably, 7M was negatively associated with perspective maintenance across conditions and predicted reduced emotional intensity during observer-perspective retrieval, consistent with a role in affective distancing or perspective instability. These findings raise the possibility that constructive memory processes are supported by partially graded functional transitions across the posterior parietal cortex—such that dorsal regions (e.g., PGa, 7A) support spatial scene construction, while more posterior or medial regions (e.g., PGp, 7P, 7M) contribute to affective and subjective dimensions of memory. While this pattern is broadly consistent with macroscale cortical gradients (Margulies et al. 2016), the spatial organization of these effects may also reflect fine-grained, interdigitated parietal networks (Braga and Buckner 2017). This framework offers a promising direction for understanding how posterior parietal systems dynamically coordinate multiple facets of experience during memory reconstruction.

Our findings contribute to constructive memory theory (Schacter and Addis 2007) by showing that visual perspective can be intentionally manipulated prior to retrieval, with measurable consequences for neural activity and subjective experience. This supports the idea that memory is not merely the reactivation of stored content, but a goal-driven process shaped by top-down cues that frame how memories are reconstructed. By isolating the preparatory and reconstructive stages of perspective-guided remembering, we provide evidence that retrieval orientation is not limited to semantic or perceptual goals (Gurguryan and Sheldon 2019) but can also be directed toward visuospatial and self-referential dimensions. Crucially, orienting toward a visual perspective modulates the mode of recollection, altering the spatial framework, self-location, and subjective vantage point, rather than merely biasing which details are retrieved. This claim is supported by our finding that parietal subregions such as PGa and PGp are engaged during the cue phase in ways that predict subsequent perspective maintenance, indicating that participants were configuring a visual vantage point prior to memory access. This preparatory recruitment, along with probe-phase effects on vividness and emotion, suggests that viewpoint-based retrieval orientation shapes the form and structure of mental simulation. This broadens the concept of retrieval orientation by showing that top-down goals can shape not just the content retrieved, but the spatial and subjective format through which a memory is mentally simulated. This extension complements prior work on semantic and perceptual goal states (Rugg and Wilding 2000) and aligns with theories of constructive episodic simulation, where scene construction, self-location, and perspective are core features of remembering (Buckner and Carroll 2007; Byrne et al. 2007; Hassabis and Maguire 2007).

Beyond theoretical implications, these findings hold important translational value. By demonstrating that visual perspective functions as a retrieval orientation, our results reframe it as an active, goal-directed control process rather than a passive feature of recollection. This reconceptualization aligns with emotion regulation frameworks and suggests new avenues for intervention. Observer perspectives are commonly used in cognitive reappraisal techniques to help individuals reinterpret aversive memories with greater psychological distance (Webb et al. 2012) and have been linked to reduced emotional reactivity in experimental settings (Kross and Ayduk 2011). However, our findings also highlight potential risks of habitual or inflexible use: overreliance on observer perspectives—particularly when engaged rigidly or by default—may impair integration and contribute to emotional disengagement, as seen in dissociative disorders (Brewin et al. 2010). While some models propose that observer perspective reflects a compensatory strategy in the aftermath of trauma (Rubin et al. 2008), our results suggest that its effectiveness may depend on the flexibility and context of its use.

Notably, activity in the 7M subregion was negatively associated with both perspective maintenance and emotional intensity during observer-perspective retrieval, suggesting a role in affective distancing or instability in viewpoint control. This region may serve as a neural marker of how individuals engage observer perspective to regulate emotional content. For example, in trauma-focused interventions, 7M activity during instructed shifts to an observer perspective could indicate whether an individual is engaging this strategy in a flexible and adaptive way. Individuals who can selectively recruit 7M in response to retrieval goals may benefit from observer-based distancing, whereas those who show blunted or habitual engagement may be at risk for emotional disengagement or fragmentation. Longitudinal research could further assess whether patterns of 7M engagement predict long-term outcomes in trauma recovery or responsiveness to memory-based therapies. These findings highlight the importance of visual perspective as a dynamic retrieval orientation, with implications for identifying neural markers of regulatory success and guiding individualized approaches to trauma memory intervention.

Interestingly, we did not observe differences in vividness or emotional intensity across retrieval perspectives. Instead, vividness and emotional intensity ratings did not significantly vary across retrieval perspective conditions when averaged across participants. This discrepancy may be attributable to a key methodological difference: in the present study, visual perspective cues were temporally separated from the AM probe. Participants had time to adopt the instructed perspective before the memory title appeared, which may have made perspective adoption less cognitively demanding compared to previous designs in which cues were presented simultaneously with the AM content (Faul et al. 2020). In such designs, participants must construct the scene and adopt a new perspective concurrently, which may amplify differences in subjective ratings due to increased demands on self-location and spatial transformation. Furthermore, prior studies that found reductions in emotional intensity when shifting to an observer perspective often deliberately selected memories strongly encoded from an own-eyes perspective, thereby requiring a true perspective shift during retrieval (St. Jacques et al. 2017). In contrast, our design matched initial visual perspective ratings across conditions, and did not require participants to shift perspectives. This likely reduced the extent of dissonance between initial encoding and instructed retrieval perspective, which may have attenuated subjective differences in emotional intensity and vividness. These findings suggest that retrieval orientation effects on subjective experience may depend on both the timing of the cue and the magnitude of perspective shift required. A promising future direction would be to directly compare memories strongly biased toward one perspective at encoding, and then manipulate whether participants are instructed to maintain or shift perspective during retrieval. Such a design would more precisely isolate the contribution of perspective transformation to changes in subjective memory experience.

While our design matched Session 1 perspective ratings across retrieval conditions, we acknowledge that, overall, memories were initially recalled from a more own-eyes-like perspective. As a result, trials in the Observer condition may have involved a greater shift from participants’ default retrieval perspective, whereas Own Eyes trials were more likely to align with the original encoding viewpoint. Importantly, both our main GLM and our brain-behavior models included Session 1 perspective bias scores, therefore capturing the degree to which each memory initially favored an own-eyes versus observer viewpoint. Including this covariate did not alter the pattern of results, supporting the interpretation that neural effects during the cue and probe phases reflect retrieval orientation and reconstructive processes, rather than differences in perspective congruence alone. Nevertheless, future work could systematically manipulate perspective congruency to more directly examine its influence on memory transformation.

### Limitations

One limitation of this study is the absence of spontaneous perspective ratings in the Retrieve condition, which precludes strong inferences about default perspective use. Future work could address this by collecting retrospective perspective ratings or incorporating trial-level assessments. Additionally, while we attempted to balance perspective biases across participants, some individuals may have a strong default preference in their adopted perspective. Pre-screening or modeling individual differences in perspective use could help clarify the stability of retrieval orientations. Finally, while our ROI analyses were grounded in cytoarchitectonic definitions from a probabilistic atlas (Eickhoff et al. 2005), the precise spatial organization of posterior parietal subregions may vary across individuals, particularly in transitional regions. This variability may limit the anatomical specificity of subregion-level interpretations (Braga and Buckner 2017).

### Conclusion

This study provides the first direct evidence that visual perspective cues initiate retrieval orientation processes that shape how AMs are later reconstructed. The AG, particularly PGp, serves as a key hub for preparing perspective-specific retrieval, while precuneus subregions contribute differentially to egocentric perspective, visual imagery, and affective modulation during memory retrieval. These findings clarify the distinct phases of perspective-guided remembering and underscore the importance of preparatory processes in shaping the subjective experience of memory.

## Data and Code Availability

Data and code supporting the findings of this study are publicly available via the Open Science Framework: https://osf.io/4785r/

## Acknowledgments

The authors thank Chloe King and Anna Cuff for their help in the preparation of fMRI scripts and the data collection. This research was completed as part of SK’s doctoral thesis at the University of Alberta. The authors used OpenAI’s ChatGPT to assist with editing and language refinement during manuscript preparation. All AI-assisted contributions were critically reviewed and revised by the authors.

## Author Contributions

SK: Software, Validation, Formal Analysis, Investigation, Data Curation, Writing – Original Draft, Visualization, Project Administration. PLS: Conceptualization, Methodology, Software, Validation, Formal Analysis, Resources, Data Curation, Writing – Review and Editing, Visualization, Supervision, Funding Acquisition. The authors declare no financial or non-financial conflict of interest.

## References

Barry DN, Maguire EA. 2019. Remote memory and the hippocampus: A constructive critique. Trends in Cognitive Sciences. 23(2):128–142. https://www.sciencedirect.com/science/article/pii/S136466131830264X.

Bergouignan L, Nyberg L, Ehrsson HH. 2014. Out-of-body-induced hippocampal amnesia. P Natl Acad Sci USA. 111(12):4421–4426. 10.1073/pnas.1318801111.

Berntsen D, Rubin DC. 2006. Emotion and vantage point in autobiographical memory. Cognition Emotion. 20(8):1193–1215. 10.1080/02699930500371190.

Berryhill ME, Phuong L, Picasso L, Cabeza R, Olson IR. 2007. Parietal lobe and episodic memory: Bilateral damage causes impaired free recall of autobiographical memory. J Neurosci. 27(52):14415–14423. 10.1523/JNEUROSCI.4163-07.2007

Binder JR, Desai RH. 2011. The neurobiology of semantic memory. Trends in Cognitive Sciences. 15(11):527–536. 10.1016/j.tics.2011.10.001.

Bonnici HM, Cheke LG, Green DAE, FitzGerald T, Simons JS. 2018. Specifying a causal role for angular gyrus in autobiographical memory. J Neurosci. 38(49):10438–10443. https://www.ncbi.nlm.nih.gov/pubmed/30355636.

Bonnici HM, Richter FR, Yazar Y, Simons JS. 2016. Multimodal feature integration in the angular gyrus during episodic and semantic retrieval. J Neurosci. 36(20):5462–5471. https://www.ncbi.nlm.nih.gov/pubmed/27194327.

Braga RM, Buckner RL. 2017. Parallel interdigitated distributed networks within the individual estimated by intrinsic functional connectivity. Neuron. 95(2):457–471.e455. 10.1016/j.neuron.2017.06.038.

Brett, M., Anton, J.-L., Valabregue, R., Poline, J. B. 2002. Region of interest analysis using an spm toolbox. 8th International Conference on Functional Mapping of the Human Brain; Sendai, Japan. Neuroimage, 16(2).

Brewin CR, Gregory JD, Lipton M, Burgess N. 2010. Intrusive images in psychological disorders: Characteristics, neural mechanisms, and treatment implications. Psychol Rev. 117(1):210–232. 10.1037/a0018113

Brunec IK, Bellana B, Ozubko JD, Man V, Robin J, Liu Z, Grady C, Rosenbaum RS, Winocur G, Barense MD et al. 2018. Multiple scales of representation along the hippocampal anteroposterior axis in humans. Curr Biol. 28(13):2129–2135.e2126. 10.1016/j.cub.2018.05.016.

Buckner RL, Carroll DC. 2007. Self-projection and the brain. Trends in Cognitive Science. 11(2):49–57. 10.1016/j.tics.2006.11.004

Butler AC, Rice HJ, Wooldridge CL, Rubin DC. 2016. Visual imagery in autobiographical memory: The role of repeated retrieval in shifting perspective. Consciousness and Cognition. 42:237–253. http://www.sciencedirect.com/science/article/pii/S1053810016300472.

Byrne P, Becker S, Burgess N. 2007. Remembering the past and imagining the future: A neural model of spatial memory and imagery. Psychol Rev. 114(2):340–375. 10.1037/0033-295X.114.2.340.

Cabeza R, St Jacques P. 2007. Functional neuroimaging of autobiographical memory. Trends Cogn Sci. 11(5):219–227. 10.1016/j.tics.2007.02.005.

Caspers S, Eickhoff SB, Geyer S, Scheperjans F, Mohlberg H, Zilles K, Amunts K. 2008. The human inferior parietal lobule in stereotaxic space. Brain Structure and Function. 212(6):481–495. 10.1007/s00429-008-0195-z.

Caspers S, Geyer S, Schleicher A, Mohlberg H, Amunts K, Zilles K. 2006. The human inferior parietal cortex: Cytoarchitectonic parcellation and interindividual variability. NeuroImage. 33(2):430–448. https://www.sciencedirect.com/science/article/pii/S1053811906006975.

Cavanna AE, Trimble MR. 2006. The precuneus: A review of its functional anatomy and behavioural correlates. Brain. 129:564–583. 10.1093/brain/awl004.

Chen G, Saad ZS, Britton JC, Pine DS, Cox RW. 2013. Linear mixed-effects modeling approach to fmri group analysis. NeuroImage. 73:176–190. https://www.sciencedirect.com/science/article/pii/S1053811913000943.

Dobbins IG, Wagner AD. 2005. Domain-general and domain-sensitive prefrontal mechanisms for recollecting events and detecting novelty. Cerebral Cortex. 15(11):1768–1778. 10.1093/cercor/bhi054

Eickhoff SB, Stephan KE, Mohlberg H, Grefkes C, Fink GR, Amunts K, Zilles K. 2005. A new spm toolbox for combining probabilistic cytoarchitectonic maps and functional imaging data. NeuroImage. 25(4):1325–1335. https://www.sciencedirect.com/science/article/pii/S105381190400792X.

Eklund A, Nichols TE, Knutsson H. 2016. Cluster failure: Why fmri inferences for spatial extent have inflated false-positive rates. Proc Natl Acad Sci U S A. 113(28):7900–7905. 10.1073/pnas.1602413113.

Faul L, St. Jacques PL, DeRosa JT, Parikh N, De Brigard F. 2020. Differential contribution of anterior and posterior midline regions during mental simulation of counterfactual and perspective shifts in autobiographical memories. NeuroImage. 215:116843. http://www.sciencedirect.com/science/article/pii/S105381192030330X.

Gilmore AW, Nelson SM, Chen H, McDermott KB. 2018. Task-related and resting-state fmri identify distinct networks that preferentially support remembering the past and imagining the future. Neuropsychologia. 110:180–189. https://www.sciencedirect.com/science/article/pii/S0028393217302282.

Grol M, Vingerhoets G, De Raedt R. 2017. Mental imagery of positive and neutral memories: A fmri study comparing field perspective imagery to observer perspective imagery. Brain and Cognition. 111:13–24. http://www.sciencedirect.com/science/article/pii/S0278262616302792.

Gurguryan L, Sheldon S. 2019. Retrieval orientation alters neural activity during autobiographical memory recollection. NeuroImage. 199:534–544. https://www.sciencedirect.com/science/article/pii/S1053811919304732.

Hassabis D, Maguire EA. 2007. Deconstructing episodic memory with construction. Trends Cogn Sci. 11(7):299–306. 10.1016/j.tics.2007.05.001

Hebscher M, Ibrahim C, Gilboa A. 2020. Precuneus stimulation alters the neural dynamics of autobiographical memory retrieval. NeuroImage. 210:116575. http://www.sciencedirect.com/science/article/pii/S1053811920300628.

Herron JE, Rugg MD. 2003. Retrieval orientation and the control of recollection. Journal of Cognitive Neuroscience. 15(6):843–854. 10.1162/jocn_a_00299

Humphreys GF, Lambon Ralph MA, Simons JS. 2021. A unifying account of angular gyrus contributions to episodic and semantic cognition. Trends in Neurosciences. 10.1016/j.tins.2021.01.006.

Iriye H, Chancel M, Ehrsson HH. 2023. Sense of own body shapes neural processes of memory encoding and reinstatement. Cerebral Cortex. 34(1). 10.1093/cercor/bhad443.

Iriye H, St Jacques PL. 2020. How visual perspective influences the spatiotemporal dynamics of autobiographical memory retrieval. Cortex. 129:464–475. https://www.ncbi.nlm.nih.gov/pubmed/32599462.

Iriye H, St. Jacques PL. 2024. An embodied perspective: Angular gyrus and precuneus decode selfhood in memories of naturalistic events. bioRxiv. 2024.2009.2009.612088. https://www.biorxiv.org/content/biorxiv/early/2024/09/10/2024.09.09.612088.full.pdf.

Jitsuishi T, Yamaguchi A. 2021. Posterior precuneus is highly connected to medial temporal lobe revealed by tractography and white matter dissection. Neuroscience. 466:173–185. 10.1016/j.neuroscience.2021.05.009.

Jitsuishi T, Yamaguchi A. 2023. Characteristic cortico-cortical connection profile of human precuneus revealed by probabilistic tractography. Scientific Reports. 13(1):1936. 10.1038/s41598-023-29251-2.

Kross E, Ayduk O. 2011. Making meaning out of negative experiences by self-distancing. Current Directions in Psychological Science. 20(3):187–191. 10.1177/0963721411408883

Maguire EA, Mullally SL. 2013. The hippocampus: A manifesto for change. Journal of Experimental Psychology: General. 142(4):1180–1189. 10.1037/a0033650

Margulies DS, Ghosh SS, Goulas A, Falkiewicz M, Huntenburg JM, Langs G, Bezgin G, Eickhoff SB, Castellanos FX, Petrides M et al. 2016. Situating the default-mode network along a principal gradient of macroscale cortical organization. Proceedings of the National Academy of Sciences. 113(44):12574–12579. https://www.pnas.org/doi/abs/10.1073/pnas.1608282113.

Margulies DS, Vincent JL, Kelly C, Lohmann G, Uddin LQ, Biswal BB, Villringer A, Castellanos FX, Milham MP, Petrides M. 2009. Precuneus shares intrinsic functional architecture in humans and monkeys. P Natl Acad Sci USA. 106(47):20069–20074. 10.1073/pnas.0905314106.

Morcom AM, Rugg MD. 2012. Retrieval orientation and the control of recollection: An fmri study. Journal of Cognitive Neuroscience. 24(12):2372–2384. 10.1162/jocn_a_00299.

Nigro G, Neisser U. 1983. Point of view in personal memories. Cognitive Psychol. 15(4):467–482. 10.1016/0010-0285(83)90016-6.

Ollinger JM, Shulman GL, Corbetta M. 2001. Separating processes within a trial in event-related functional mri: i. The method. Neuroimage. 13(1):210–217. 10.1006/nimg.2000.0710.

Peirce J, Gray JR, Simpson S, MacAskill M, Höchenberger R, Sogo H, Kastman E, Lindeløv JK. 2019. Psychopy2: Experiments in behavior made easy. Behavior Research Methods. 51(1):195–203. 10.3758/s13428-018-01193-y.

Poppenk J, Evensmoen HR, Moscovitch M, Nadel L. 2013. Long-axis specialization of the human hippocampus. Trends in Cognitive Sciences. 17(5):230–240. 10.1016/j.tics.2013.03.005.

Rice HJ, Rubin DC. 2009. I can see it both ways: First- and third-person visual perspectives at retrieval. Consciousness and Cognition. 18(4):877–890. 10.1016/j.concog.2009.07.004.

Ritchey M, Cooper RA. 2020. Deconstructing the posterior medial episodic network. Trends Cogn Sci. 24(6):451–465. https://www.ncbi.nlm.nih.gov/pubmed/32340798.

Ritchey M, Montchal ME, Yonelinas AP, Ranganath C. 2015. Delay-dependent contributions of medial temporal lobe regions to episodic memory retrieval. eLife. 4:e05025. 10.7554/eLife.05025.

Rubin DC, Berntsen D, Bohni MK. 2008. A memory-based model of posttraumatic stress disorder: Evaluating basic assumptions underlying the ptsd diagnosis. Psychol Rev. 115(4):985–1011. https://www.ncbi.nlm.nih.gov/pubmed/18954211.

Rubin DC, Umanath S. 2015. Event memory: A theory of memory for laboratory, autobiographical, and fictional events. Psychol Rev. 122(1):1–23. 10.1037/a0037907

Rugg MD, Wilding EL. 2000. Retrieval processing and episodic memory. Trends Cogn Sci. 4(3):108–115. 10.1016/S1364-6613(00)01445-5

Schacter DL, Addis DR. 2007. The cognitive neuroscience of constructive memory: Remembering the past and imagining the future. Philos Trans R Soc Lond B Biol Sci. 362(1481):773–786. https://www.ncbi.nlm.nih.gov/pubmed/17395575.

Scheperjans F, Hermann K, Eickhoff SB, Amunts K, Schleicher A, Zilles K. 2007. Observer-independent cytoarchitectonic mapping of the human superior parietal cortex. Cerebral Cortex. 18(4):846–867. 10.1093/cercor/bhm116.

Seghier ML. 2013. The angular gyrus: Multiple functions and multiple subdivisions. Neuroscientist. 19(1):43–61. https://www.ncbi.nlm.nih.gov/pubmed/22547530.

St. Jacques, PL. 2025. Visual Perspective Shapes Subjective Experience: Dissociable Parietal Contributions to the Constructive Nature of Memory. Retrieved from osf.io/preprints/psyarxiv/pmgb7_v2

St. Jacques PL. 2019. A new perspective on visual perspective in memory. Current Directions in Psychological Science. 28(5):450–455. https://journals.sagepub.com/doi/abs/10.1177/0963721419850158.

St. Jacques PL. 2022. Visual perspective in event memory. In: Brady TM, Brainbridge WA, editors. Visual memory. Routledge. p. 278–297.

St Jacques PL, Olm C, Schacter DL. 2013. Neural mechanisms of reactivation-induced updating that enhance and distort memory. P Natl Acad Sci USA. 110(49):19671–19678. 10.1073/pnas.1319630110.

St Jacques PL, Szpunar KK, Schacter DL. 2017. Shifting visual perspective during retrieval shapes autobiographical memories. Neuroimage. 148(1):103–114. https://www.ncbi.nlm.nih.gov/pubmed/27989780.

St. Jacques PL, Carpenter AC, Szpunar KK, Schacter DL. 2018. Remembering and imagining alternative versions of the personal past. Neuropsychologia. 110:170–179. https://www.sciencedirect.com/science/article/pii/S0028393217302270.

St. Jacques PL, Kragel PA, Rubin DC. 2013. Neural networks supporting autobiographical memory retrieval in posttraumatic stress disorder. Cognitive, Affective, & Behavioral Neuroscience. 13(3):554–566. 10.3758/s13415-013-0157-7.

St. Jacques PL, Szpunar KK, Schacter DL. 2017. Shifting visual perspective during retrieval shapes autobiographical memories. NeuroImage. 148:103–114. 10.1016/j.neuroimage.2016.12.028

Stark CEL, Squire LR. 2001. When zero is not zero: The problem of ambiguous baseline conditions in fmri. Proceedings of the National Academy of Sciences. 98(22):12760–12766. https://www.pnas.org/doi/abs/10.1073/pnas.221462998.

Webb TL, Miles E, Sheeran P. 2012. Dealing with feeling: A meta-analysis of the effectiveness of strategies derived from the process model of emotion regulation. Psychological Bulletin. 138(4):775–808. 10.1037/a0027600

Yazar Y, Bergström ZM, Simons JS. 2017. Reduced multimodal integration of memory features following continuous theta burst stimulation of angular gyrus. Brain Stimulation. 10(3):624–629. https://www.sciencedirect.com/science/article/pii/S1935861X17306186

