## Supplemental Table 1 for "Functional Specialization of Angular Gyrus and Precuneus Subregions for Perspective-Guided Autobiographical Memory Retrieval"

Accepted at *Imaging Neuroscience*

**Supplemental Results: Brain-Behavior Relationship with Categorical ROI Comparison**

We conducted exploratory linear mixed-effects models in which ROI was treated as a categorical factor (PGa vs PGp; 7A vs 7M vs 7P) testing Region × fMRI signal × Condition interactions (see Supplemental Tables 17 – 28).

***Vividness***. During the cue phase, there was a significant condition and percent signal change (fMRI activity) interaction in the AG (see Supplemental Table 17), specifically when comparing the Own Eyes and Observer conditions ($\beta$ = -.143, *SE* = .053, 95% CI [-.25, -.04], *t* (4152.7) = -2.68, *p* = .007). To follow up on the significant interaction, we estimated simple slopes of percent signal change on vividness within each retrieval orientation condition. A significant negative slope was observed in the Observer ($\beta$ = -.134, SE = .037, 95% CI [-.21, -.06], t (4148) = -3.63, p < .001) and Retrieve conditions ($\beta$ = -.078, SE = .037, 95% CI [-.15, -.01], t (4151) = -2.11, p = .035), indicating that greater AG recruitment was associated with reduced vividness. The corresponding slope in the Own Eyes condition ($\beta$ = .008, SE = .039, 95% CI [-.07, .08], t (4144) = .21, p = .834) was not statistically significant. In contrast, there were no interactions between percent signal change when comparing the Own Eyes versus Retrieve ($\beta$ = -.086, SE = .053, 95% CI [-.19, .02], t (4142.8) = -1.63, p = .103) or the Observer versus Retrieve conditions ($\beta$ = .056, SE = .052, 95% CI [-.05, .15], t (4140.6) = 1.08, p = .280).

During the probe phase, there was a significant condition and percent signal change (fMRI activity) interaction in the AG (see Supplemental Table 18), specifically when comparing the Own Eyes and Observer conditions ($\beta$ = .146, *SE* = .055, 95% CI [.04, .26], *t* (4032.5) = 2.64, *p* = .008). To follow up on the significant interaction, we estimated simple slopes of percent signal change on vividness within each retrieval orientation condition. A significant positive slope was observed in the Observer condition ($\beta$ = .165, SE = .047, 95% CI [.07, .26], t (87.8) = 3.51, p < .001), indicating that greater AG recruitment was associated with higher vividness. The corresponding slope in the Own Eyes ($\beta$ = .018, SE = .047, 95% CI [-.08, .11], t (89.6) = .40, p = .693) and Retrieve conditions ($\beta$ = .091, SE = .047, 95% CI [-.00, .19], t (99.6) = 1.93, p = .057) were not statistically significant. In contrast, there were no interactions between percent signal change when comparing the Own Eyes versus Retrieve ($\beta$ = .072, SE = .056, 95% CI [-.04, .18], t (4112.9) = 1.29, p = .198) or the Observer versus Retrieve conditions ($\beta$ = -.073, SE = .056, 95% CI [-.18, .04], t (4119.6) = -1.33, p = .189).

Turning to the precuneus, during the cue phase (see Supplemental Table 19) there was a significant condition and percent signal change (fMRI activity) interaction, specifically when comparing the Own Eyes and Retrieve conditions ($\beta$ = -.096, *SE* = .035, 95% CI [-.17, .03], *t* (6214.9) = -2.70, *p* = .007). To follow up on the significant interaction, we estimated simple slopes of percent signal change on vividness within each retrieval orientation condition. However, simple effects revealed no significant effect in any of the retrieval orientation conditions (Own Eyes: $\beta$ = .057, SE = .035, 95% CI [-.01, .13], t (59.9) = 1.61, p = .113; Observer: $\beta$ = -.002, SE = .035, 95% CI [-.07, .07], t (58.5) = -.06, p = .954; Retrieve: $\beta$ = -.039, SE = .035, 95% CI [-.11, .03], t (60.2) = -1.10, p = .278). There were no interactions between percent signal change when comparing the Own Eyes versus Observer ($\beta$ = .072, SE = .056, 95% CI [-.04, .18], t (4112.9) = 1.29, p = .198) or the Observer versus Retrieve conditions ($\beta$ = -.037, SE = .035, 95% CI [-.11, .03], t (6212.9) = -1.05, p = .292).

During the probe phase, retrieval orientation condition was a significant predictor of vividness ratings in precuneus (see Supplemental Table 20), when comparing the Own Eyes and

Retrieve ($\beta$ = .065, SE = .024, 95% CI [.02, .11], t(6212.7) = 2.74, p = .006), and Observer and Retrieve conditions ($\beta$ = .101, SE = .024, 95% CI [.05, .15], t(6218.9) = 4.20, p < .001). Additionally, percent signal change during the cue phase was a significant predictor of vividness in Session 2 ($\beta$ = .056, SE = .016, 95% CI [.02, .09], t(6276.3) = 3.43, p < .001). However, precuneus subregion, or two-way interactions between variables did not significantly predict vividness ratings in Session 2 (all *p*s > .140).

***Emotional Intensity***. During the cue phase, there were no significant differences in the AG (all *p*s > .187; see Supplemental Table 21). During the probe phase (see Supplemental Table 22), percent signal change in AG significantly predicted emotional intensity ratings ($\beta$ = .103, SE = .048, 95% CI [.01, .20], t (35.8) = 2.13, p = .198), indicating increased neural recruitment in AG during the probe phase is associated with higher emotional intensity. There were no other significant predictors or interactions.

Turning to the precuneus, during the cue phase there was a significant condition and percent signal change (fMRI activity) interaction (see Supplemental Table 23), specifically when comparing the Own Eyes and Retrieve ($\beta$ = .090, *SE* = .036, 95% CI [.02, .16], *t* (5937.3) = 2.51, *p* = .012) and Observer and Retrieve conditions ($\beta$ = .091, *SE* = .036, 95% CI [.02, .16], *t* (5984.4) = 2.54, *p* = .011). To follow up on the significant interaction, we estimated simple slopes of percent signal change on emotional intensity within each retrieval orientation condition. A significant negative slope was observed in the Retrieve condition ($\beta$ = -.073, SE = .030, 95% CI [-.13, -.01], t (123) = -2.47, p = .015), indicating that greater precuneus recruitment was associated with lower vividness. The corresponding slope in the Own Eyes ($\beta$ = .017, SE = .029, 95% CI [-.04, .08], t (119) = .59, p = .558) and Observer conditions ($\beta$ = .018, SE = .029, 95% CI [-

.04, .08], t (117) = .62, p = .535) were not statistically significant. In contrast, there were no interactions between percent signal change when comparing the Own Eyes versus Observer conditions (*β* = .001, SE = .036, 95% CI [-.07, .07], t (5806.3) = .03, p = .977).

During the probe phase, within the precuneus there was a significant condition and percent signal change (fMRI activity) interaction (see Supplemental Table 24), specifically when comparing the Observer and Retrieve conditions (*β* = -.107, *SE* = .035, 95% CI [-.18, -.04], *t* (6135.1) = -3.02, *p* = .003). To follow up on the significant interaction, we estimated simple slopes of percent signal change on emotional intensity within each retrieval orientation condition. However, simple effects revealed no significant effect in any of the retrieval orientation conditions (Own Eyes: *β* = -.012, SE = .037, 95% CI [-.09, .06], t (55.1) = -.34, p = .733; Observer: *β* = -.070, SE = .037, 95% CI [-.15, .01], t (57.7) = -1.87, p = .067; Retrieve: *β* = .037, SE = .037, 95% CI [-.04, .11], t (56.8) = .98, p = .329). There were no interactions between percent signal change when comparing the Own Eyes versus Retrieve (*β* = -.049, SE = .035, 95% CI [-.12, .02], t (6120.9) = -1.41, p = .158) or the Own Eyes versus Observer conditions (*β* = -.057, SE = .035, 95% CI [-.13, .01], t (6136.1) = -1.64, p = .101).

***Perspective Maintenance***. During the cue phase, there was a significant condition and percent signal change (fMRI activity) interaction in the AG (see Supplemental Table 25), specifically when comparing the Own Eyes and Observer (*β* = -.141, *SE* = .067, 95% CI [-.27, -.01], *t* (4117.9) = -2.09, *p* = .037), and Own Eyes and Retrieve conditions (*β* = -.162, *SE* = .067, 95% CI [-.29, -.03], *t* (4110.7) = -2.41, *p* = .016). To follow up on the significant interaction, we estimated simple slopes of percent signal change on perspective maintenance within each retrieval orientation condition. A significant positive slope was observed in the Own Eyes condition ($\beta$ = .120, SE = .049, 95% CI [.02, .22], t (4106) = 2.43, p = .015), indicating that greater AG recruitment was associated with higher perspective maintenance. The corresponding slope in the Observer ($\beta$ = -.020, SE = .047, 95% CI [-.11, .07], t (4115) = -.44, p = .662) and Retrieve conditions ($\beta$ = -.041, SE = .047, 95% CI [-.13, .05], t (4120) = -.88, p = .378) were not statistically significant. In contrast, there were no interactions between percent signal change when comparing the Observer versus Retrieve conditions ($\beta$ = -.020, SE = .065, 95% CI [-.15, .11], t (4110.4) = -.32, p = .752).

During the probe phase in the AG (see Supplemental Table 26), retrieval orientation was a significant predictor of perspective maintenance ratings, when comparing the Own Eyes and Observer ($\beta$ = -.329, SE = .053, 95% CI [-.43, -.22], t(4067.8) = -6.16, p = .038), Own Eyes and Retrieve ($\beta$ = .440, SE = .053, 95% CI [.01, .22], t(4070.2) = 2.07, p < .001), and Observer and Retrieve conditions ($\beta$ = .111, SE = .053, 95% CI [.34, .55], t(4068) = 8.26, p < .001). AG subregion, percent signal change during the probe phase, or two-way interactions between variables did not significantly predict perspective maintenance ratings in Session 2 (all *p*s > .163).

Turning to the precuneus, during the cue phase there was a significant condition and percent signal change (fMRI activity) interaction (see Supplemental Table 27), specifically when comparing the Own Eyes and Observer conditions ($\beta$ = -.120, *SE* = .045, 95% CI [-.21, -.03], *t* (6097.4) = -2.67, *p* = .008). To follow up on the significant interaction, we estimated simple slopes of percent signal change on perspective maintenance within each retrieval orientation condition. A significant positive slope was observed in the Own Eyes condition ($\beta$ = .106, SE = .039, 95% CI [.03, .19], t (76.9) = 2.66, p = .009), indicating that greater precuneus recruitment was associated with higher perspective maintenance. The corresponding slope in the Observer ($\beta$ = -.014, SE = .039, 95% CI [-.09, .07], t (76.9) = -.35, p = .727) and Retrieve conditions ($\beta$ = .039, SE = .039, 95% CI [-.04, .12], t (77) = 1, p = .321) were not statistically significant. In contrast, there were no interactions between percent signal change when comparing the Own Eyes versus Retrieve ($\beta$ = -.066, SE = .045, 95% CI [-.15, .02], t (6147.3) = -1.48, p = .140) and Observer versus Retrieve conditions ($\beta$ = .053, SE = .044, 95% CI [-.03, .14], t (6166.3) = 1.20, p = .229).

During the probe phase within the precuneus (see Supplemental Table 28), there was a significant three-way interaction between precuneus subregion, condition, and percent signal change (fMRI activity), specifically when comparing the 7A and 7M activity in the Own Eyes versus Observer ($\beta$ = .280, *SE* = .119, 95% CI [.05, .51], *t* (6135.8) = 2.35, *p* = .019), and in the Own Eyes versus Retrieve conditions ($\beta$ = .266, *SE* = .119, 95% CI [.03, .50], *t* (6133.3) = 2.23, *p* = .026). To follow up on the significant interaction, we estimated simple slopes of percent signal change on perspective maintenance in each retrieval orientation condition for each precuneus subregion. Only significant slope was observed in the Own Eyes condition for 7A ($\beta$ = .238, SE = .078, 95% CI [.08, .39], t (769) = 3.05, p = .002) and 7P activity ($\beta$ = .118, SE = .059, 95% CI [.00, .23], t (257) = 2, p = .047), indicating that greater 7A and 7P recruitment was associated with higher perspective maintenance. The corresponding slopes in other conditions for each region were not significant (all *p*s > .124).

### Supplemental Tables

**Supplemental Table 1. Model Predicting Vividness from Left AG (Cue Phase)**

| Fixed Effect | Estimate (β) | SE | 95% CI Lower | 95% CI Upper | t-value | p-value |
|---|---|---|---|---|---|---|
| Intercept | 3.531 | 0.072 | 3.389 | 3.672 | 48.92 | <.001 |
| Retrieval Condition: Own Eyes | 0.0316 | 0.042 | -0.051 | 0.114 | 0.748 | .455 |
| Retrieval Condition: Retrieve | 0.094 | 0.042 | 0.011 | 0.177 | 2.22 | .026 |
| Percent Signal Change (PGa) | -0.015 | 0.040 | -0.092 | 0.062 | -0.39 | .696 |
| Percent Signal Change (PGp) | -0.065 | 0.046 | -0.156 | 0.025 | -1.41 | .159 |
| PGa x Own Eyes | 0.182 | 0.071 | 0.042 | 0.322 | 2.561 | 0.010 |
| PGa x Retrieve | 0.072 | 0.069 | -0.064 | 0.209 | 1.039 | 0.299 |
| Vividness (S1 covariate) | 0.246 | 0.029 | 0.189 | 0.302 | 8.47 | <.001 |
| Emotion (S1 covariate) | 0.134 | 0.018 | 0.099 | 0.170 | 7.51 | <.001 |
| Perspective Bias (S1 covariate) | 0.040 | 0.014 | 0.011 | 0.069 | 2.72 | .013 |

Note. The reference level for the retrieval orientation factor is ‘Observer’. PGa = percent signal change in left anterior angular gyrus during the cue phase. PGp = percent signal change in left posterior angular gyrus during the cue phase. Session 1 ratings were included as covariates to account for baseline variation in subjective memory experience.

**Supplemental Table 2. Model Predicting Vividness from Precuneus (Cue Phase)**

| Fixed Effect | Estimate (β) | SE | 95% CI Lower | 95% CI Upper | t-value | p-value |
|---|---|---|---|---|---|---|
| Intercept | 3.529 | 0.072 | 3.387 | 3.671 | 48.76 | <.001 |
| Retrieval Condition: Own Eyes | -0.068 | 0.042 | -0.151 | 0.015 | -1.60 | .109 |
| Retrieval Condition: Observer | -0.099 | 0.042 | -0.182 | -0.016 | -2.33 | .020 |
| Percent Signal Change (7A) | -0.041 | 0.058 | -0.157 | 0.073 | -0.71 | .477 |
| Percent Signal Change (7M) | -0.023 | 0.034 | -0.090 | 0.044 | -0.67 | .502 |
| Percent Signal Change (7P) | 0.035 | 0.049 | -0.062 | 0.132 | 0.70 | .482 |
| 7A x Own Eyes | 0.102 | 0.083 | -0.059 | 0.265 | 1.24 | .216 |
| 7A x Observer | 0.028 | 0.084 | -0.136 | 0.193 | 0.34 | .738 |
| Vividness (S1 covariate) | 0.244 | 0.029 | 0.187 | 0.301 | 8.41 | <.001 |
| Emotion (S1 covariate) | 0.135 | 0.018 | 0.100 | 0.171 | 7.55 | <.001 |
| Perspective Bias (S1 covariate) | 0.040 | 0.015 | 0.011 | 0.069 | 2.75 | .012 |

Note. The reference level for the retrieval orientation factor is ‘Retrieve’. 7A = percent signal change in 7A during the cue phase. 7M = percent signal change in 7M during the cue phase. 7P = percent signal change in 7P during the cue phase. Session 1 ratings were included as covariates to account for baseline variation in subjective memory experience.

**Supplemental Table 3. Model Predicting Vividness from Left AG (Probe Phase)**

| Fixed Effect | Estimate (β) | SE | 95% CI Lower | 95% CI Upper | t-value | p-value |
|---|---|---|---|---|---|---|
| Intercept | 3.526 | 0.073 | 3.382 | 3.670 | 48.14 | <.001 |
| Retrieval Condition: Own Eyes | -0.074 | 0.042 | -0.157 | 0.009 | -1.75 | .081 |
| Retrieval Condition: Observer | -0.117 | 0.042 | -0.201 | -0.034 | -2.76 | .006 |
| Percent Signal Change (PGa) | -0.014 | 0.043 | -0.100 | 0.071 | -0.33 | .744 |
| Percent Signal Change (PGp) | 0.109 | 0.049 | 0.012 | 0.206 | 2.22 | .027 |
| PGp x Own Eyes | -0.057 | 0.083 | -0.220 | 0.105 | -0.69 | .490 |
| PGp x Observer | 0.079 | 0.081 | -0.080 | 0.238 | 0.98 | .330 |
| Vividness (S1 covariate) | 0.244 | 0.028 | 0.188 | 0.301 | 8.49 | <.001 |
| Emotion (S1 covariate) | 0.135 | 0.018 | 0.100 | 0.170 | 7.51 | <.001 |
| Perspective Bias (S1 covariate) | 0.041 | 0.014 | 0.012 | 0.070 | 2.79 | .011 |

Note. The reference level for the retrieval orientation factor is ‘Retrieve’. PGa = percent signal change in left anterior angular gyrus during the probe phase. PGp = percent signal change in left posterior angular gyrus during the probe phase. Session 1 ratings were included as covariates to account for baseline variation in subjective memory experience.

**Supplemental Table 4. Model Predicting Vividness from Precuneus (Probe Phase)**

| Fixed Effect | Estimate (β) | SE | 95% CI Lower | 95% CI Upper | t-value | p-value |
|---|---|---|---|---|---|---|
| Intercept | 3.529 | 0.072 | 3.389 | 3.669 | 49.39 | <.001 |
| Retrieval Condition: Own Eyes | -0.075 | 0.042 | -0.158 | 0.007 | -1.77 | .076 |
| Retrieval Condition: Observer | -0.115 | 0.043 | -0.199 | -0.032 | -2.72 | .007 |
| Percent Signal Change (7A) | -0.029 | 0.063 | -0.152 | 0.094 | -0.46 | .643 |
| Percent Signal Change (7M) | -0.038 | 0.032 | -0.101 | 0.025 | -1.17 | .243 |
| Percent Signal Change (7P) | 0.132 | 0.052 | 0.029 | 0.234 | 2.53 | .011 |
| 7P x Own Eyes | -0.032 | 0.060 | -0.150 | 0.085 | -0.54 | .591 |
| 7P x Observer | 0.020 | 0.060 | -0.098 | 0.140 | 0.34 | .731 |
| Vividness (S1 covariate) | 0.240 | 0.028 | 0.184 | 0.297 | 8.37 | <.001 |
| Emotion (S1 covariate) | 0.137 | 0.018 | 0.100 | 0.172 | 7.63 | <.001 |
| Perspective Bias (S1 covariate) | 0.041 | 0.014 | 0.012 | 0.069 | 2.84 | .010 |

Note. The reference level for the retrieval orientation factor is ‘Retrieve’. 7A = percent signal change in 7A during the cue phase. 7M = percent signal change in 7M during the cue phase. 7P = percent signal change in 7P during the cue phase. Session 1 ratings were included as covariates to account for baseline variation in subjective memory experience.

**Supplemental Table 5. Model Predicting Emotional Intensity from Left AG (Cue Phase)**

| Fixed Effect | Estimate (β) | SE | 95% CI Lower | 95% CI Upper | t-value | p-value |
|---|---|---|---|---|---|---|
| Intercept | 3.326 | 0.091 | 3.148 | 3.505 | 36.46 | < .001 |
| Retrieval Condition: Own Eyes | -0.046 | 0.043 | -0.131 | 0.038 | -1.07 | 0.284 |
| Retrieval Condition: Observer | -0.090 | 0.043 | -0.175 | -0.005 | -2.09 | 0.036 |
| Percent Signal Change (PGa) | -0.062 | 0.040 | -0.142 | 0.017 | -1.54 | 0.124 |
| Percent Signal Change (PGp) | 0.057 | 0.047 | -0.035 | 0.150 | 1.22 | 0.224 |
| PGa x Own Eyes | 0.088 | 0.073 | -0.054 | 0.233 | 1.22 | 0.222 |
| PGa x Observer | 0.018 | 0.071 | -0.121 | 0.158 | 0.26 | 0.796 |
| Vividness (S1 covariate) | 0.054 | 0.029 | -0.002 | 0.111 | 1.87 | 0.072 |
| Emotion (S1 covariate) | 0.438 | 0.033 | 0.372 | 0.504 | 12.95 | < .001 |
| Perspective Bias (S1 covariate) | -0.001 | 0.012 | -0.024 | 0.022 | -0.10 | 0.926 |

Note. The reference level for the retrieval orientation factor is ‘Retrieve’. PGa = percent signal change in left anterior angular gyrus during the cue phase. PGp = percent signal change in left posterior angular gyrus during the cue phase. Session 1 ratings were included as covariates to account for baseline variation in subjective memory experience.

**Supplemental Table 6. Model Predicting Emotional Intensity from Precuneus (Cue Phase)**

| Fixed Effect | Estimate (β) | SE | 95% CI Lower | 95% CI Upper | t-value | p-value |
|---|---|---|---|---|---|---|
| Intercept | 3.325 | 0.090 | 3.147 | 3.502 | 36.65 | < .001 |
| Retrieval Condition: Own Eyes | -0.038 | 0.043 | -0.123 | 0.045 | -0.90 | 0.369 |
| Retrieval Condition: Observer | -0.082 | 0.043 | -0.166 | 0.002 | -1.90 | 0.058 |
| Percent Signal Change (7A) | -0.039 | 0.060 | -0.157 | 0.078 | -0.66 | 0.509 |
| Percent Signal Change (7M) | -0.042 | 0.034 | -0.111 | 0.025 | -1.22 | 0.221 |
| Percent Signal Change (7P) | 0.048 | 0.051 | -0.051 | 0.148 | 0.95 | 0.342 |
| 7A x Own Eyes | 0.100 | 0.102 | -0.099 | 0.300 | 0.99 | 0.323 |
| 7A x Observer | 0.127 | 0.107 | -0.083 | 0.337 | 1.18 | 0.236 |
| 7M x Own Eyes | 0.045 | 0.067 | -0.086 | 0.178 | 0.68 | 0.499 |
| 7M x Observer | 0.031 | 0.069 | -0.104 | 0.166 | 0.45 | 0.655 |
| Vividness (S1 covariate) | 0.052 | 0.028 | -0.003 | 0.109 | 1.83 | 0.078 |
| Emotion (S1 covariate) | 0.439 | 0.033 | 0.373 | 0.506 | 12.96 | < .001 |
| Perspective Bias (S1 covariate) | 0 | 0.012 | -0.024 | 0.020 | -0.07 | 0.946 |

Note. The reference level for the retrieval orientation factor is ‘Retrieve’. 7A = percent signal change in 7A during the cue phase. 7M = percent signal change in 7M during the cue phase. 7P = percent signal change in 7P during the cue phase. Session 1 ratings were included as covariates to account for baseline variation in subjective memory experience.

**Supplemental Table 7. Model Predicting Emotional Intensity from Left AG (Probe Phase)**

| Fixed Effect | Estimate (β) | SE | 95% CI Lower | 95% CI Upper | t-value | p-value |
|---|---|---|---|---|---|---|
| Intercept | 3.324 | 0.091 | 3.145 | 3.502 | 36.47 | < .001 |
| Retrieval Condition: Own Eyes | -0.043 | 0.043 | -0.128 | 0.040 | -1.01 | 0.311 |
| Retrieval Condition: Observer | -0.094 | 0.043 | -0.179 | -0.009 | -2.17 | 0.030 |
| Percent Signal Change (PGa) | 0.077 | 0.044 | -0.009 | 0.165 | 1.74 | 0.081 |
| Percent Signal Change (PGp) | 0.027 | 0.050 | -0.072 | 0.126 | 0.54 | 0.591 |
| PGp x Own Eyes | -0.068 | 0.084 | -0.233 | 0.097 | -0.81 | 0.420 |
| PGp x Observer | -0.007 | 0.082 | -0.169 | 0.154 | -0.09 | 0.930 |
| Vividness (S1 covariate) | 0.054 | 0.029 | -0.002 | 0.111 | 1.86 | 0.072 |
| Emotion (S1 covariate) | 0.436 | 0.033 | 0.370 | 0.503 | 12.93 | < .001 |
| Perspective Bias (S1 covariate) | 0.00 | 0.012 | -0.023 | 0.024 | 0.03 | 0.977 |

Note. The reference level for the retrieval orientation factor is 'Retrieve'. PGa = percent signal change in left anterior angular gyrus during the probe phase. PGp = percent signal change in left posterior angular gyrus during the probe phase. Session 1 ratings were included as covariates to account for baseline variation in subjective memory experience.

**Supplemental Table 8. Model Predicting Emotional Intensity from Precuneus (Probe Phase)**

| Fixed Effect | Estimate (β) | SE | 95% CI Lower | 95% CI Upper | t-value | p-value |
|---|---|---|---|---|---|---|
| Intercept | 3.326 | 0.090 | 3.148 | 3.503 | 36.73 | < .001 |
| Retrieval Condition: Own Eyes | -0.043 | 0.043 | -0.127 | 0.040 | -1.01 | 0.311 |
| Retrieval Condition: Retrieve | -0.086 | 0.043 | -0.171 | -0.001 | -1.99 | 0.046 |
| Percent Signal Change (7A) | -0.038 | 0.063 | -0.163 | 0.086 | -0.607 | 0.544 |
| Percent Signal Change (7M) | -0.084 | 0.033 | -0.149 | -0.018 | -2.527 | 0.012 |
| Percent Signal Change (7P) | 0.069 | 0.053 | -0.035 | 0.174 | 1.298 | 0.194 |
| 7M x Own Eyes | -0.03 | 0.05 | -0.13 | 0.07 | -0.54 | 0.587 |
| 7M x Retrieve | -0.12 | 0.05 | -0.22 | -0.01 | -2.23 | 0.026 |
| Vividness (S1 covariate) | 0.05 | 0.03 | 0 | 0.11 | 1.87 | 0.072 |
| Emotion (S1 covariate) | 0.44 | 0.03 | 0.37 | 0.51 | 13.04 | < .001 |
| Perspective Bias (S1 covariate) | 0 | 0.01 | -0.02 | 0.02 | -0.04 | 0.971 |

Note. The reference level for the retrieval orientation factor is 'Observer'. 7A = percent signal change in 7A during the cue phase. 7M = percent signal change in 7M during the cue phase. 7P = percent signal change in 7P during the cue phase. Session 1 ratings were included as covariates to account for baseline variation in subjective memory experience.

**Supplemental Table 9. Model Predicting Perspective Maintenance from Left AG (Cue Phase)**

| Fixed Effect | Estimate (β) | SE | 95% CI Lower | 95% CI Upper | t-value | p-value |
|---|---|---|---|---|---|---|
| Intercept | 3.664 | 0.086 | 3.495 | 3.383 | 42.53 | <.001 |
| Retrieval Condition: Observer | -0.334 | 0.053 | -0.439 | -0.229 | -6.25 | <.001 |
| Retrieval Condition: Retrieve | 0.113 | 0.053 | 0.008 | 0.218 | 2.11 | .035 |
| Percent Signal Change (PGa) | 0.098 | 0.050 | 0 | 0.197 | 1.97 | .049 |
| Percent Signal Change (PGp) | -0.088 | 0.058 | -0.203 | 0.026 | -1.51 | .131 |
| PGa x Observer | -0.209 | 0.090 | -0.386 | -0.031 | -2.31 | 0.021 |
| PGa x Retrieve | -0.238 | 0.090 | -0.415 | -0.060 | -2.63 | .009 |
| Vividness (S1 covariate) | 0.074 | 0.025 | 0.024 | 0.125 | 2.90 | .004 |
| Emotion (S1 covariate) | 0.061 | 0.022 | 0.017 | 0.105 | 2.71 | .007 |
| Perspective Bias (S1 covariate) | 0.038 | 0.018 | 0.003 | 0.074 | 2.14 | .043 |

Note. The reference level for the retrieval orientation factor is 'Own Eyes'. PGa = percent signal change in left anterior angular gyrus during the probe phase. PGp = percent signal change in left posterior angular gyrus during the probe phase. Session 1 ratings were included as covariates to account for baseline variation in subjective memory experience.

**Supplemental Table 10. Model Predicting Perspective Maintenance from Precuneus (Cue Phase)**

| Fixed Effect | Estimate (β) | SE | 95% CI Lower | 95% CI Upper | t-value | p-value |
|---|---|---|---|---|---|---|
| Intercept | 3.660 | 0.087 | 3.489 | 3.831 | 41.90 | <.001 |
| Retrieval Condition: Observer | -0.327 | 0.053 | -0.432 | -0.222 | -6.10 | <.001 |
| Retrieval Condition: Retrieve | 0.125 | 0.053 | 0.021 | 0.230 | 2.35 | .019 |
| Percent Signal Change (7A) | 0.167 | 0.073 | 0.023 | 0.312 | 2.27 | .023 |
| Percent Signal Change (7M) | -0.020 | 0.043 | -0.105 | 0.063 | -0.49 | .627 |
| Percent Signal Change (7P) | -0.058 | 0.062 | -0.180 | 0.064 | -0.93 | .352 |
| 7A x Observer | -0.224 | 0.104 | -0.429 | -0.019 | -2.14 | .032 |
| 7A x Retrieve | -0.214 | 0.103 | -0.417 | -0.010 | -2.06 | .039 |
| Vividness (S1 covariate) | 0.073 | 0.025 | 0.022 | 0.123 | 2.83 | .005 |
| Emotion (S1 covariate) | 0.063 | 0.022 | 0.019 | 0.107 | 2.81 | .005 |
| Perspective Bias (S1 covariate) | 0.039 | 0.018 | 0.003 | 0.075 | 2.14 | .042 |

Note. The reference level for the retrieval orientation factor is 'Own Eyes'. 7A = percent signal change in 7A during the cue phase. 7M = percent signal change in 7M during the cue phase. 7P = percent signal change in 7P during the cue phase. Session 1 ratings were included as covariates to account for baseline variation in subjective memory experience.

**Supplemental Table 11. Model Predicting Perspective Maintenance from Left AG (Probe Phase)**

| Fixed Effect | Estimate (β) | SE | 95% CI Lower | 95% CI Upper | t-value | p-value |
|---|---|---|---|---|---|---|
| Intercept | 3.657 | 0.084 | 3.491 | 3.823 | 43.14 | <.001 |
| Retrieval Condition: Own Eyes | 0.340 | 0.053 | 0.234 | 0.443 | 6.31 | <.001 |
| Retrieval Condition: Retrieve | 0.470 | 0.054 | 0.364 | 0.576 | 8.71 | <.001 |
| Percent Signal Change (PGa) | -0.095 | 0.055 | -0.203 | 0.013 | -1.72 | .086 |
| Percent Signal Change (PGp) | 0.145 | 0.062 | 0.023 | 0.267 | 2.33 | .020 |
| PGp x Own Eyes | -0.099 | 0.102 | -0.299 | 0.101 | -0.07 | .332 |
| PGp x Retrieve | -0.105 | 0.102 | -0.307 | 0.095 | -1.03 | .302 |
| Vividness (S1 covariate) | 0.071 | 0.025 | 0.020 | 0.122 | 2.77 | .006 |
| Emotion (S1 covariate) | 0.064 | 0.022 | 0.020 | 0.109 | 2.86 | .004 |
| Perspective Bias (S1 covariate) | 0.040 | 0.018 | 0.004 | 0.075 | 2.22 | .035 |

Note. The reference level for the retrieval orientation factor is ‘Observer’. PGa = percent signal change in left anterior angular gyrus during the probe phase. PGp = percent signal change in left posterior angular gyrus during the probe phase. Session 1 ratings were included as covariates to account for baseline variation in subjective memory experience.

**Supplemental Table 12. Model Predicting Perspective Maintenance from Precuneus (Probe Phase: Interaction Model)**

| Fixed Effect | Estimate (β) | SE | 95% CI Lower | 95% CI Upper | t-value | p-value |
|---|---|---|---|---|---|---|
| Intercept | 3.658 | 0.081 | 3.498 | 3.818 | 44.81 | <.001 |
| Retrieval Condition: Observer | -0.331 | 0.053 | -0.436 | -0.226 | -6.19 | <.001 |
| Retrieval Condition: Retrieve | 0.130 | 0.053 | 0.026 | 0.235 | 2.45 | .014 |
| Percent Signal Change (7A) | 0.061 | 0.079 | -0.094 | 0.216 | 0.77 | .442 |
| Percent Signal Change (7M) | -0.183 | 0.040 | -0.263 | -0.103 | -4.48 | <.001 |
| Percent Signal Change (7P) | 0.151 | 0.065 | 0.022 | 0.280 | 2.30 | .022 |
| 7A x Observer | -0.410 | 0.167 | -0.738 | -0.081 | -2.44 | .014 |
| 7A x Retrieve | -0.333 | 0.161 | -0.649 | -0.017 | -2.07 | .015 |
| 7P x Observer | 0.144 | 0.123 | -0.098 | 0.386 | 1.17 | 0.244 |
| 7P x Retrieve | 0.036 | 0.116 | -0.191 | 0.264 | 0.32 | 0.753 |
| Vividness (S1 covariate) | 0.063 | 0.025 | 0.013 | 0.114 | 2.47 | .014 |
| Emotion (S1 covariate) | 0.068 | 0.022 | 0.024 | 0.112 | 3.05 | .002 |
| Perspective Bias (S1 covariate) | 0.040 | 0.018 | 0.005 | 0.076 | 2.24 | .035 |

Note. The reference level for the retrieval orientation factor is ‘Own Eyes’. 7A = percent signal change in 7A during the cue phase. 7M = percent signal change in 7M during the cue phase. 7P = percent signal change in 7P during the cue phase. Session 1 ratings were included as covariates to account for baseline variation in subjective memory experience.

**Supplemental Table 13. Model Predicting Vividness from AG (Cue Phase: Categorical Approach for AG Subregions)**

| Fixed Effect | Estimate (β) | SE | 95% CI Lower | 95% CI Upper | t-value | p-value |
|---|---|---|---|---|---|---|
| Intercept | 3.539 | 0.074 | 3.394 | 3.685 | 47.66 | <.001 |
| Retrieval Condition: Observer | -0.352 | 0.041 | -0.116 | 0.046 | -0.84 | .399 |
| Retrieval Condition: Retrieve | 0.063 | 0.041 | -0.018 | 0.144 | 1.51 | .131 |
| AG Subregion[1] | 0.003 | 0.024 | -0.043 | 0.050 | 0.15 | .878 |
| % Sig. Change[2] | -0.057 | 0.028 | -0.114 | -0.001 | -2.01 | .044 |
| AG Subregion x Condition[3] | 0.013 | 0.059 | -0.102 | 0.128 | 0.22 | .823 |
| AG Subregion x Condition[4] | 0.000 | 0.059 | -0.114 | 0.116 | 0.01 | .989 |
| Condition x % Sig. Change[5] | -0.143 | 0.053 | -0.247 | -0.038 | -2.68 | .007 |
| Condition x % Sig. Change[6] | -0.086 | 0.053 | -0.191 | 0.017 | -1.63 | .103 |
| AG Subregion x % Sig. Change | -0.021 | 0.043 | -0.106 | 0.063 | -0.49 | .621 |
| Vividness (S1 covariate) | 0.235 | 0.030 | 0.176 | 0.294 | 7.84 | <.001 |
| Emotion (S1 covariate) | 0.132 | 0.021 | 0.091 | 0.174 | 6.23 | <.001 |
| Perspective Bias (S1 covariate) | 0.038 | 0.015 | 0.008 | 0.068 | 2.55 | .018 |

Note. The reference level for the retrieval orientation factor is 'Own Eyes'. Session 1 ratings were included as covariates to account for baseline variation in subjective memory experience.

[1] PGa vs. PGp

[2] Percent signal change in left AG during the cue phase

[3] Interaction between left AG subregions and Condition (Own Eyes vs. Observer)

[4] Interaction between left AG subregions and Condition (Own Eyes vs. Retrieve)

[5] Interaction between Condition (Own Eyes vs. Observer) and percent signal change

[6] Interaction between Condition (Own Eyes vs. Retrieve) and percent signal change

**Supplemental Table 14. Model Predicting Vividness from AG (Probe Phase: Categorical Approach for AG Subregions)**

| Fixed Effect | Estimate (β) | SE | 95% CI Lower | 95% CI Upper | t-value | p-value |
|---|---|---|---|---|---|---|
| Intercept | 3.550 | 0.075 | 3.402 | 3.698 | 47.10 | <.001 |
| Retrieval Condition: Observer | -0.019 | 0.042 | -0.101 | 0.063 | -0.45 | .650 |
| Retrieval Condition: Retrieve | 0.073 | 0.042 | -0.009 | 0.156 | 1.74 | .082 |
| AG Subregion[1] | -0.022 | 0.025 | -0.071 | 0.026 | -0.89 | .373 |
| % Sig. Change[2] | 0.070 | 0.040 | -0.009 | 0.149 | 1.72 | .091 |
| AG Subregion x Condition[3] | -0.046 | 0.060 | -0.165 | 0.072 | -0.77 | .440 |
| AG Subregion x Condition[4] | -0.015 | 0.060 | -0.132 | 0.103 | -0.25 | .804 |
| Condition x % Sig. Change[5] | 0.146 | 0.055 | 0.037 | 0.255 | 2.64 | .008 |
| Condition x % Sig. Change[6] | 0.072 | 0.056 | -0.038 | 0.184 | 1.29 | .198 |
| AG Subregion x % Sig. Change | 0.044 | 0.047 | -0.049 | 0.137 | 0.92 | .356 |
| Vividness (S1 covariate) | 0.233 | 0.029 | 0.175 | 0.291 | 7.87 | <.001 |
| Emotion (S1 covariate) | 0.129 | 0.021 | 0.088 | 0.170 | 6.16 | <.001 |
| Perspective Bias (S1 covariate) | 0.039 | 0.015 | 0.009 | 0.069 | 2.54 | .018 |

Note. The reference level for the retrieval orientation factor is ‘Own Eyes’. Session 1 ratings were included as covariates to account for baseline variation in subjective memory experience.

[1] PGa vs. PGp

[2] Percent signal change in left AG during the probe phase

[3] Interaction between left AG subregions and Condition (Own Eyes vs. Observer)

[4] Interaction between left AG subregions and Condition (Own Eyes vs. Retrieve)

[5] Interaction between Condition (Own Eyes vs. Observer) and percent signal change

[6] Interaction between Condition (Own Eyes vs. Retrieve) and percent signal change

**Supplemental Table 15. Model Predicting Vividness from Precuneus (Cue Phase: Categorical Approach for Precuneus Subregions)**

| Fixed Effect | Estimate (β) | SE | 95% CI Lower | 95% CI Upper | t-value | p-value |
|---|---|---|---|---|---|---|
| Intercept | 3.530 | 0.078 | 3.376 | 3.684 | 44.89 | <.001 |
| Retrieval Condition: Observer | -0.034 | 0.041 | -0.115 | 0.046 | -0.83 | .404 |
| Retrieval Condition: Retrieve | 0.059 | 0.041 | -0.021 | 0.140 | 1.45 | .148 |
| Precuneus Subregion: 7M | -0.017 | 0.042 | -0.100 | 0.065 | -0.41 | .680 |
| Precuneus Subregion: 7P | 0.005 | 0.041 | -0.075 | 0.086 | 0.14 | .887 |
| % Sig. Change[1] | 0.048 | 0.046 | -0.042 | 0.139 | 1.05 | .296 |
| Precuneus Subregion x Condition[2] | 0.008 | 0.059 | -0.107 | 0.124 | 0.14 | .889 |
| Precuneus Subregion x Condition[3] | 0.017 | 0.059 | -0.098 | 0.133 | 0.30 | .765 |
| Precuneus Subregion x Condition[4] | -0.004 | 0.058 | -0.118 | 0.109 | -0.08 | .939 |
| Precuneus Subregion x Condition[5] | -0.006 | 0.058 | -0.121 | 0.107 | -0.12 | .906 |
| Condition x % Sig. Change[6] | -0.059 | 0.035 | -0.129 | 0.009 | -1.68 | .093 |
| Condition x % Sig. Change[7] | -0.096 | 0.035 | -0.167 | -0.026 | -2.70 | .007 |
| Precuneus Subregion x % Sig. Change[8] | 0.014 | 0.040 | -0.064 | 0.093 | 0.36 | .719 |
| Precuneus Subregion x % Sig. Change[9] | 0.012 | 0.041 | -0.068 | 0.092 | 0.30 | .762 |
| Vividness (S1 covariate) | 0.234 | 0.029 | 0.176 | 0.292 | 7.93 | <.001 |
| Emotion (S1 covariate) | 0.131 | 0.021 | 0.088 | 0.174 | 6.02 | <.001 |
| Perspective Bias (S1 covariate) | 0.028 | 0.015 | 0.008 | 0.068 | 2.53 | .018 |

Note. The reference level for the retrieval orientation factor is ‘Own Eyes’. The reference level for the precuneus subregion factor is ‘7A’. Session 1 ratings were included as covariates to account for baseline variation in subjective memory experience.

[1] Percent signal change in the precuneus during the cue phase

[2] Interaction between precuneus subregions (7A vs. 7M) and Condition (Own Eyes vs. Observer)

[3] Interaction between precuneus subregions (7A vs. 7M) and Condition (Own Eyes vs. Retrieve)

[4] Interaction between precuneus subregions (7A vs. 7P) and Condition (Own Eyes vs. Observer)

[5] Interaction between precuneus subregions (7A vs. 7P) and Condition (Own Eyes vs. Retrieve)

[6] Interaction between Condition (Own Eyes vs. Observer) and percent signal change

[7] Interaction between Condition (Own Eyes vs. Retrieve) and percent signal change

[8] Interaction between precuneus subregions (7A vs. 7M) and percent signal change

[9] Interaction between precuneus subregions (7A vs. 7P) and percent signal change

**Supplemental Table 16. Model Predicting Vividness from Precuneus (Probe Phase: Categorical Approach for Precuneus Subregions)**

| Fixed Effect | Estimate (β) | SE | 95% CI Lower | 95% CI Upper | t-value | p-value |
|---|---|---|---|---|---|---|
| Intercept | 3.547 | 0.073 | 3.404 | 3.690 | 48.52 | <.001 |
| Retrieval Condition: Observer | -0.035 | 0.024 | -0.082 | 0.011 | -1.47 | .141 |
| Retrieval Condition: Retrieve | 0.065 | 0.024 | 0.018 | 0.112 | 2.74 | .006 |
| Precuneus Subregion: 7M | -0.018 | 0.025 | -0.067 | 0.030 | -0.73 | .463 |
| Precuneus Subregion: 7P | -0.007 | 0.024 | -0.055 | 0.041 | -0.29 | .769 |
| % Sig. Change[1] | 0.056 | 0.016 | 0.024 | 0.089 | 3.43 | <.001 |
| Precuneus Subregion x Condition[2] | -0.003 | 0.060 | -0.121 | 0.114 | -0.05 | .956 |
| Precuneus Subregion x Condition[3] | 0.004 | 0.060 | -0.113 | 0.122 | 0.07 | .942 |
| Precuneus Subregion x Condition[4] | -0.004 | 0.058 | -0.119 | 0.110 | -0.08 | .935 |
| Precuneus Subregion x Condition[5] | 0.003 | 0.058 | -0.111 | 0.118 | 0.06 | .950 |
| Condition x % Sig. Change[6] | 0.009 | 0.034 | -0.058 | 0.077 | 0.27 | .784 |
| Condition x % Sig. Change[7] | -0.013 | 0.034 | -0.081 | 0.054 | -0.40 | .692 |
| Precuneus Subregion x % Sig. Change[8] | -0.037 | 0.039 | -0.115 | 0.040 | -0.95 | .343 |
| Precuneus Subregion x % Sig. Change[9] | 0.004 | 0.041 | -0.075 | 0.084 | 0.11 | .911 |
| Vividness (S1 covariate) | 0.231 | 0.029 | 0.173 | 0.290 | 7.77 | <.001 |
| Emotion (S1 covariate) | 0.132 | 0.021 | 0.089 | 0.175 | 6.08 | <.001 |
| Perspective Bias (S1 covariate) | 0.037 | 0.015 | 0.008 | 0.067 | 2.49 | .020 |

Note. The reference level for the retrieval orientation factor is ‘Own Eyes’. The reference level for the precuneus subregion factor is ‘7A’. Session 1 ratings were included as covariates to account for baseline variation in subjective memory experience.

[1] Percent signal change in the precuneus during the probe phase

[2] Interaction between precuneus subregions (7A vs. 7M) and Condition (Own Eyes vs. Observer)

[3] Interaction between precuneus subregions (7A vs. 7M) and Condition (Own Eyes vs. Retrieve)

[4] Interaction between precuneus subregions (7A vs. 7P) and Condition (Own Eyes vs. Observer)

[5] Interaction between precuneus subregions (7A vs. 7P) and Condition (Own Eyes vs. Retrieve)

[6] Interaction between Condition (Own Eyes vs. Observer) and percent signal change

[7] Interaction between Condition (Own Eyes vs. Retrieve) and percent signal change

[8] Interaction between precuneus subregions (7A vs. 7M) and percent signal change

[9] Interaction between precuneus subregions (7A vs. 7P) and percent signal change

**Supplemental Table 17. Model Predicting Emotional Intensity from AG (Cue Phase: Categorical Approach for AG Subregions)**

| Fixed Effect | Estimate (β) | SE | 95% CI Lower | 95% CI Upper | t-value | p-value |
|---|---|---|---|---|---|---|
| Intercept | 3.340 | 0.093 | 3.158 | 3.523 | 35.87 | <.001 |
| Retrieval Condition: Observer | -0.045 | 0.042 | -0.128 | 0.037 | -1.07 | .287 |
| Retrieval Condition: Retrieve | 0.035 | 0.042 | -0.047 | 0.118 | 0.84 | .400 |
| AG Subregion[1] | -0.000 | 0.024 | -0.048 | 0.047 | -0.03 | .974 |
| % Sig. Change[2] | -0.025 | 0.029 | -0.083 | 0.031 | -0.88 | .380 |
| AG Subregion x Condition[3] | 0.002 | 0.060 | -0.115 | 0.120 | 0.44 | .965 |
| AG Subregion x Condition[4] | 0.003 | 0.059 | -0.113 | 0.121 | 0.59 | .953 |
| Condition x % Sig. Change[5] | -0.031 | 0.054 | -0.137 | 0.074 | -0.58 | .560 |
| Condition x % Sig. Change[6] | -0.071 | 0.054 | -0.177 | 0.034 | -1.32 | .188 |
| AG Subregion x % Sig. Change | 0.036 | 0.044 | -0.049 | 0.123 | 0.83 | .407 |
| Vividness (S1 covariate) | 0.061 | 0.027 | 0.007 | 0.114 | 2.23 | .034 |
| Emotion (S1 covariate) | 0.426 | 0.033 | 0.361 | 0.491 | 12.88 | <.001 |
| Perspective Bias (S1 covariate) | -0.006 | 0.016 | -0.037 | 0.025 | -0.38 | .707 |

Note. The reference level for the retrieval orientation factor is 'Own Eyes'. Session 1 ratings were included as covariates to account for baseline variation in subjective memory experience.

[1] PGa vs. PGp

[2] Percent signal change in left AG during the cue phase

[3] Interaction between left AG subregions and Condition (Own Eyes vs. Observer)

[4] Interaction between left AG subregions and Condition (Own Eyes vs. Retrieve)

[5] Interaction between Condition (Own Eyes vs. Observer) and percent signal change

[6] Interaction between Condition (Own Eyes vs. Retrieve) and percent signal change

**Supplemental Table 18. Model Predicting Emotional Intensity from AG (Probe Phase: Categorical Approach for AG Subregions)**

| Fixed Effect | Estimate (β) | SE | 95% CI Lower | 95% CI Upper | t-value | p-value |
|---|---|---|---|---|---|---|
| Intercept | 3.351 | 0.092 | 3.168 | 3.533 | 36.06 | <.001 |
| Retrieval Condition: Observer | -0.055 | 0.042 | -0.138 | 0.028 | -1.29 | .197 |
| Retrieval Condition: Retrieve | 0.046 | 0.042 | -0.037 | 0.130 | 1.08 | .280 |
| AG Subregion[1] | -0.027 | 0.025 | -0.077 | 0.023 | -1.06 | .290 |
| % Sig. Change[2] | 0.103 | 0.048 | 0.008 | 0.198 | 2.13 | .040 |
| AG Subregion x Condition[3] | 0.003 | 0.061 | -0.117 | 0.123 | 0.05 | .960 |
| AG Subregion x Condition[4] | -0.007 | 0.060 | -0.126 | 0.111 | -0.12 | .901 |
| Condition x % Sig. Change[5] | -0.005 | 0.056 | -0.116 | 0.105 | -0.09 | .926 |
| Condition x % Sig. Change[6] | 0.051 | 0.057 | -0.059 | 0.163 | 0.90 | .364 |
| AG Subregion x % Sig. Change | -0.018 | 0.048 | -0.113 | 0.077 | -0.38 | .707 |
| Vividness (S1 covariate) | 0.059 | 0.027 | 0.005 | 0.113 | 2.17 | .038 |
| Emotion (S1 covariate) | 0.425 | 0.032 | 0.361 | 0.489 | 13.02 | <.001 |
| Perspective Bias (S1 covariate) | -0.004 | 0.016 | -0.037 | 0.028 | -0.26 | .801 |

Note. The reference level for the retrieval orientation factor is ‘Own Eyes’. Session 1 ratings were included as covariates to account for baseline variation in subjective memory experience.

[1] PGa vs. PGp

[2] Percent signal change in left AG during the probe phase

[3] Interaction between left AG subregions and Condition (Own Eyes vs. Observer)

[4] Interaction between left AG subregions and Condition (Own Eyes vs. Retrieve)

[5] Interaction between Condition (Own Eyes vs. Observer) and percent signal change

[6] Interaction between Condition (Own Eyes vs. Retrieve) and percent signal change

**Supplemental Table 19. Model Predicting Emotional Intensity from Precuneus (Cue Phase: Categorical Approach for Precuneus Subregions)**

| Fixed Effect | Estimate (β) | SE | 95% CI Lower | 95% CI Upper | t-value | p-value |
|---|---|---|---|---|---|---|
| Intercept | 3.345 | 0.092 | 3.163 | 3.526 | 36.12 | <.001 |
| Retrieval Condition: Own Eyes | -0.035 | 0.024 | -0.083 | 0.012 | -1.46 | .144 |
| Retrieval Condition: Observer | -0.079 | 0.024 | -0.127 | -0.032 | -3.27 | .001 |
| Precuneus Subregion: 7M | 0.004 | 0.024 | -0.044 | 0.053 | 0.18 | .855 |
| Precuneus Subregion: 7P | -0.000 | 0.024 | -0.048 | 0.047 | -0.01 | .995 |
| % Sig. Change[1] | -0.012 | 0.021 | -0.053 | 0.028 | -0.59 | .556 |
| Precuneus Subregion x Condition[2] | -0.018 | 0.060 | -0.136 | 0.099 | -0.31 | .760 |
| Precuneus Subregion x Condition[3] | -0.022 | 0.060 | -0.140 | 0.094 | -0.38 | .705 |
| Precuneus Subregion x Condition[4] | 0.003 | 0.059 | -0.113 | 0.119 | 0.05 | .958 |
| Precuneus Subregion x Condition[5] | 0.002 | 0.059 | -0.113 | 0.119 | 0.05 | .962 |
| Condition x % Sig. Change[6] | 0.090 | 0.036 | 0.019 | 0.161 | 2.51 | .012 |
| Condition x % Sig. Change[7] | 0.091 | 0.036 | 0.021 | 0.162 | 2.54 | .011 |
| Precuneus Subregion x % Sig. Change[8] | 0.006 | 0.040 | -0.073 | 0.085 | 0.16 | .874 |
| Precuneus Subregion x % Sig. Change[9] | 0.015 | 0.041 | -0.065 | 0.097 | 0.38 | .704 |
| Vividness (S1 covariate) | 0.063 | 0.026 | 0.012 | 0.115 | 2.42 | .022 |
| Emotion (S1 covariate) | 0.422 | 0.032 | 0.358 | 0.486 | 12.95 | <.001 |
| Perspective Bias (S1 covariate) | -0.009 | 0.017 | -0.044 | 0.025 | -0.55 | .586 |

Note. The reference level for the retrieval orientation factor is 'Retrieve'. The reference level for the precuneus subregion factor is '7A'. Session 1 ratings were included as covariates to account for baseline variation in subjective memory experience.

[1] Percent signal change in the precuneus during the cue phase

[2] Interaction between precuneus subregions (7A vs. 7M) and Condition (Retrieve vs. Own Eyes)

[3] Interaction between precuneus subregions (7A vs. 7M) and Condition (Retrieve vs. Observer)

[4] Interaction between precuneus subregions (7A vs. 7P) and Condition (Retrieve vs. Own Eyes)

[5] Interaction between precuneus subregions (7A vs. 7P) and Condition (Retrieve vs. Observer)

[6] Interaction between Condition (Retrieve vs. Own Eyes) and percent signal change

[7] Interaction between Condition (Retrieve vs. Observer) and percent signal change

[8] Interaction between precuneus subregions (7A vs. 7M) and percent signal change

[9] Interaction between precuneus subregions (7A vs. 7P) and percent signal change

**Supplemental Table 20. Model Predicting Emotional Intensity from Precuneus (Probe Phase: Categorical Approach for Precuneus Subregions)**

| Fixed Effect | Estimate (β) | SE | 95% CI Lower | 95% CI Upper | t-value | p-value |
|---|---|---|---|---|---|---|
| Intercept | 3.348 | 0.092 | 3.166 | 3.530 | 36.14 | <.001 |
| Retrieval Condition: Own Eyes | -0.034 | 0.024 | -0.081 | 0.013 | -1.40 | .161 |
| Retrieval Condition: Observer | -0.076 | 0.024 | -0.123 | -0.028 | -3.11 | .002 |
| Precuneus Subregion: 7M | 0.003 | 0.025 | -0.047 | 0.053 | 0.13 | .896 |
| Precuneus Subregion: 7P | 0.001 | 0.025 | -0.047 | 0.051 | 0.08 | .936 |
| % Sig. Change[1] | -0.015 | 0.031 | -0.077 | 0.046 | -0.49 | .629 |
| Precuneus Subregion x Condition[2] | 0.017 | 0.060 | -0.101 | 0.136 | 0.29 | .772 |
| Precuneus Subregion x Condition[3] | 0.044 | 0.060 | -0.074 | 0.164 | 0.73 | .463 |
| Precuneus Subregion x Condition[4] | 0.002 | 0.059 | -0.113 | 0.118 | 0.05 | .964 |
| Precuneus Subregion x Condition[5] | 0.013 | 0.059 | -0.102 | 0.130 | 0.23 | .819 |
| Condition x % Sig. Change[6] | -0.049 | 0.035 | -0.119 | 0.019 | -1.41 | .158 |
| Condition x % Sig. Change[7] | -0.107 | 0.035 | -0.177 | -0.037 | -3.02 | .003 |
| Precuneus Subregion x % Sig. Change[8] | -0.001 | 0.040 | -0.081 | 0.078 | -0.04 | .966 |
| Precuneus Subregion x % Sig. Change[9] | -0.004 | 0.041 | -0.086 | 0.077 | -0.11 | .911 |
| Vividness (S1 covariate) | 0.063 | 0.026 | 0.012 | 0.115 | 2.43 | .022 |
| Emotion (S1 covariate) | 0.422 | 0.032 | 0.358 | 0.486 | 12.86 | <.001 |
| Perspective Bias (S1 covariate) | -0.007 | 0.016 | -0.040 | 0.025 | -0.46 | .648 |

Note. The reference level for the retrieval orientation factor is 'Retrieve'. The reference level for the precuneus subregion factor is '7A'. Session 1 ratings were included as covariates to account for baseline variation in subjective memory experience.

[1] Percent signal change in the precuneus during the probe phase
[2] Interaction between precuneus subregions (7A vs. 7M) and Condition (Retrieve vs. Own Eyes)
[3] Interaction between precuneus subregions (7A vs. 7M) and Condition (Retrieve vs. Observer)
[4] Interaction between precuneus subregions (7A vs. 7P) and Condition (Retrieve vs. Own Eyes)
[5] Interaction between precuneus subregions (7A vs. 7P) and Condition (Retrieve vs. Observer)
[6] Interaction between Condition (Retrieve vs. Own Eyes) and percent signal change
[7] Interaction between Condition (Retrieve vs. Observer) and percent signal change
[8] Interaction between precuneus subregions (7A vs. 7M) and percent signal change
[9] Interaction between precuneus subregions (7A vs. 7P) and percent signal change

**Supplemental Table 21. Model Predicting Perspective Maintenance from AG (Cue Phase: Categorical Approach for AG Subregions)**

| Fixed Effect | Estimate (β) | SE | 95% CI Lower | 95% CI Upper | t-value | p-value |
|---|---|---|---|---|---|---|
| Intercept | 3.683 | 0.090 | 3.507 | 3.860 | 40.87 | <.001 |
| Retrieval Condition: Observer | -0.332 | 0.052 | -0.436 | -0.229 | -6.29 | <.001 |
| Retrieval Condition: Retrieve | 0.116 | 0.052 | 0.012 | 0.219 | 2.21 | .027 |
| AG Subregion[1] | -0.003 | 0.030 | -0.063 | 0.055 | -0.13 | .900 |
| % Sig. Change[2] | 0.045 | 0.036 | -0.026 | 0.116 | 1.24 | .215 |
| AG Subregion x Condition[3] | 0.015 | 0.074 | -0.131 | 0.162 | 0.21 | .834 |
| AG Subregion x Condition[4] | 0.012 | 0.074 | -0.133 | 0.158 | 0.17 | .864 |
| Condition x % Sig. Change[5] | -0.141 | 0.067 | -0.273 | -0.008 | -2.09 | .037 |
| Condition x % Sig. Change[6] | -0.162 | 0.067 | -0.294 | -0.030 | -2.41 | .016 |
| AG Subregion x % Sig. Change | -0.051 | 0.054 | -0.159 | 0.056 | -0.94 | .349 |
| Vividness (S1 covariate) | 0.064 | 0.026 | 0.011 | 0.117 | 2.39 | .024 |
| Emotion (S1 covariate) | 0.056 | 0.024 | 0.009 | 0.104 | 2.35 | .027 |
| Perspective Bias (S1 covariate) | 0.033 | 0.018 | -0.001 | 0.069 | 1.86 | .077 |

Note. The reference level for the retrieval orientation factor is 'Own Eyes'. Session 1 ratings were included as covariates to account for baseline variation in subjective memory experience.

[1] PGa vs. PGp

[2] Percent signal change in left AG during the cue phase

[3] Interaction between left AG subregions and Condition (Own Eyes vs. Observer)

[4] Interaction between left AG subregions and Condition (Own Eyes vs. Retrieve)

[5] Interaction between Condition (Own Eyes vs. Observer) and percent signal change

[6] Interaction between Condition (Own Eyes vs. Retrieve) and percent signal change

**Supplemental Table 22. Model Predicting Perspective Maintenance from AG (Probe Phase: Categorical Approach for AG Subregions)**

| Fixed Effect | Estimate (β) | SE | 95% CI Lower | 95% CI Upper | t-value | p-value |
|---|---|---|---|---|---|---|
| Intercept | 3.689 | 0.088 | 3.514 | 3.863 | 41.50 | <.001 |
| Retrieval Condition: Observer | -0.329 | 0.053 | -0.434 | -0.224 | -6.16 | <.001 |
| Retrieval Condition: Retrieve | 0.111 | 0.053 | 0.006 | 0.216 | 2.07 | .038 |
| AG Subregion[1] | 0.000 | 0.031 | -0.061 | 0.062 | 0.02 | .983 |
| % Sig. Change[2] | -0.004 | 0.052 | -0.107 | 0.099 | -0.08 | .940 |
| AG Subregion x Condition[3] | 0.002 | 0.077 | -0.148 | 0.153 | 0.03 | .974 |
| AG Subregion x Condition[4] | 0.022 | 0.076 | -0.126 | 0.171 | 0.30 | .764 |
| Condition x % Sig. Change[5] | -0.026 | 0.070 | -0.164 | 0.111 | -0.38 | .704 |
| Condition x % Sig. Change[6] | -0.077 | 0.070 | -0.215 | 0.061 | -1.09 | .277 |
| AG Subregion x % Sig. Change | 0.083 | 0.059 | -0.033 | 0.200 | 1.39 | .164 |
| Vividness (S1 covariate) | 0.066 | 0.026 | 0.014 | 0.118 | 2.49 | .020 |
| Emotion (S1 covariate) | 0.054 | 0.024 | 0.008 | 0.102 | 2.29 | .030 |
| Perspective Bias (S1 covariate) | 0.028 | 0.018 | -0.007 | 0.064 | 1.57 | .131 |

Note. The reference level for the retrieval orientation factor is ‘Own Eyes’. Session 1 ratings were included as covariates to account for baseline variation in subjective memory experience.

[1] PGa vs. PGp

[2] Percent signal change in left AG during the probe phase

[3] Interaction between left AG subregions and Condition (Own Eyes vs. Observer)

[4] Interaction between left AG subregions and Condition (Own Eyes vs. Retrieve)

[5] Interaction between Condition (Own Eyes vs. Observer) and percent signal change

[6] Interaction between Condition (Own Eyes vs. Retrieve) and percent signal change

**Supplemental Table 23. Model Predicting Perspective Maintenance from Precuneus (Cue Phase: Categorical Approach for Precuneus Subregions)**

| Fixed Effect | Estimate (β) | SE | 95% CI Lower | 95% CI Upper | t-value | p-value |
|---|---|---|---|---|---|---|
| Intercept | 3.686 | 0.090 | 3.509 | 3.864 | 40.76 | <.001 |
| Retrieval Condition: Own Eyes | -0.322 | 0.030 | -0.381 | -0.262 | -10.57 | <.001 |
| Retrieval Condition: Observer | 0.126 | 0.030 | 0.067 | 0.186 | 4.18 | <.001 |
| Precuneus Subregion: 7M | -0.013 | 0.031 | -0.074 | 0.046 | -0.45 | .654 |
| Precuneus Subregion: 7P | -0.004 | 0.030 | -0.063 | 0.055 | -0.13 | .893 |
| % Sig. Change[1] | 0.044 | 0.030 | -0.015 | 0.103 | 1.45 | .158 |
| Precuneus Subregion x Condition[2] | 0.023 | 0.075 | -0.123 | 0.170 | 0.31 | .755 |
| Precuneus Subregion x Condition[3] | 0.015 | 0.074 | -0.130 | 0.162 | 0.21 | .832 |
| Precuneus Subregion x Condition[4] | -0.006 | 0.074 | -0.151 | 0.139 | -0.08 | .935 |
| Precuneus Subregion x Condition[5] | -0.001 | 0.073 | -0.146 | 0.142 | -0.03 | .979 |
| Condition x % Sig. Change[6] | -0.120 | 0.045 | -0.208 | -0.032 | -2.67 | .008 |
| Condition x % Sig. Change[7] | -0.066 | 0.045 | -0.154 | 0.021 | -1.48 | .140 |
| Precuneus Subregion x % Sig. Change[8] | -0.076 | 0.050 | -0.175 | 0.023 | -1.51 | .132 |
| Precuneus Subregion x % Sig. Change[9] | -0.066 | 0.051 | -0.168 | 0.035 | -1.28 | .200 |
| Vividness (S1 covariate) | 0.058 | 0.027 | 0.005 | 0.111 | 2.16 | .040 |
| Emotion (S1 covariate) | 0.057 | 0.025 | 0.007 | 0.106 | 2.27 | .032 |
| Perspective Bias (S1 covariate) | 0.029 | 0.018 | -0.006 | 0.065 | 1.59 | .127 |

Note. The reference level for the retrieval orientation factor is ‘Own Eyes’. The reference level for the precuneus subregion factor is ‘7A’. Session 1 ratings were included as covariates to account for baseline variation in subjective memory experience.

[1] Percent signal change in the precuneus during the cue phase
[2] Interaction between precuneus subregions (7A vs. 7M) and Condition (Own Eyes vs. Observer)
[3] Interaction between precuneus subregions (7A vs. 7M) and Condition (Own Eyes vs. Retrieve)
[4] Interaction between precuneus subregions (7A vs. 7P) and Condition (Own Eyes vs. Observer)
[5] Interaction between precuneus subregions (7A vs. 7P) and Condition (Own Eyes vs. Retrieve)
[6] Interaction between Condition (Own Eyes vs. Observer) and percent signal change
[7] Interaction between Condition (Own Eyes vs. Retrieve) and percent signal change
[8] Interaction between precuneus subregions (7A vs. 7M) and percent signal change
[9] Interaction between precuneus subregions (7A vs. 7P) and percent signal change

**Supplemental Table 24. Model Predicting Perspective Maintenance from Precuneus (Probe Phase: Categorical Approach for Precuneus Subregions)**

| Fixed Effect | Estimate (β) | SE | 95% CI Lower | 95% CI Upper | t-value | p-value |
|---|---|---|---|---|---|---|
| Intercept | 3.697 | 0.091 | 3.518 | 3.877 | 40.41 | <.001 |
| Retrieval Condition: Own Eyes | -0.365 | 0.054 | -0.472 | -0.257 | -6.66 | <.001 |
| Retrieval Condition: Observer | 0.069 | 0.055 | 0.038 | 0.178 | 1.26 | .207 |
| Precuneus Subregion: 7M | -0.006 | 0.032 | -0.069 | 0.056 | -0.21 | .834 |
| Precuneus Subregion: 7P | -0.011 | 0.031 | -0.072 | 0.049 | -0.37 | .708 |
| % Sig. Change[1] | 0.054 | 0.050 | -0.043 | 0.152 | 1.10 | .275 |
| Precuneus Subregion x Condition[2] | 0.035 | 0.077 | -0.116 | 0.186 | 0.45 | .651 |
| Precuneus Subregion x Condition[3] | 0.035 | 0.075 | -0.112 | 0.184 | 0.47 | .635 |
| Precuneus Subregion x Condition[4] | 0.064 | 0.077 | -0.086 | 0.216 | 0.84 | .401 |
| Precuneus Subregion x Condition[5] | 0.042 | 0.076 | -0.107 | 0.192 | 0.56 | .577 |
| Condition x % Sig. Change[6] | -0.239 | 0.101 | -0.438 | -0.040 | -2.36 | .018 |
| Condition x % Sig. Change[7] | -0.312 | 0.102 | -0.513 | -0.110 | -3.04 | .002 |
| Precuneus Subregion x % Sig. Change[8] | -0.091 | 0.050 | -0.191 | 0.007 | -1.80 | .071 |
| Precuneus Subregion x % Sig. Change[9] | -0.014 | 0.052 | -0.116 | 0.087 | -0.28 | .779 |
| Precuneus Subregion x % Sig. Change x Condition[10] | 0.280 | 0.119 | 0.047 | 0.514 | 2.35 | .019 |
| Precuneus Subregion x % Sig. Change x Condition[11] | 0.154 | 0.125 | -0.090 | 0.399 | 1.24 | .217 |
| Precuneus Subregion x % Sig. Change x Condition[12] | 0.266 | 0.119 | 0.032 | 0.501 | 2.23 | .026 |
| Precuneus Subregion x % Sig. Change x Condition[13] | 0.162 | 0.126 | -0.084 | 0.409 | 1.29 | .197 |
| Vividness (S1 covariate) | 0.059 | 0.027 | 0.006 | 0.112 | 2.19 | .038 |
| Emotion (S1 covariate) | 0.058 | 0.024 | 0.010 | 0.106 | 2.39 | .024 |
| Perspective Bias (S1 covariate) | 0.028 | 0.019 | -0.008 | 0.066 | 1.52 | .144 |

Note. The reference level for the retrieval orientation factor is 'Own Eyes'. The reference level for the precuneus subregion factor is '7A'. Session 1 ratings were included as covariates to account for baseline variation in subjective memory experience.

[1] Percent signal change in the precuneus during the probe phase

[2] Interaction between precuneus subregions (7A vs. 7M) and Condition (Own Eyes vs. Observer)

[3] Interaction between precuneus subregions (7A vs. 7M) and Condition (Own Eyes vs. Retrieve)

[4] Interaction between precuneus subregions (7A vs. 7P) and Condition (Own Eyes vs. Observer)

[5] Interaction between precuneus subregions (7A vs. 7P) and Condition (Own Eyes vs. Retrieve)

[6] Interaction between Condition (Own Eyes vs. Observer) and percent signal change
[7] Interaction between Condition (Own Eyes vs. Retrieve) and percent signal change
[8] Interaction between precuneus subregions (7A vs. 7M) and percent signal change
[9] Interaction between precuneus subregions (7A vs. 7P) and percent signal change
[10] Three-way interactions between precuneus subregions (7A vs.7M), percent signal change, and condition (Own Eyes vs. Observer)
[11] Three-way interactions between precuneus subregions (7A vs.7P), percent signal change, and condition (Own Eyes vs. Observer)
[12] Three-way interactions between precuneus subregions (7A vs.7M), percent signal change, and condition (Own Eyes vs. Retrieve)
[13] Three-way interactions between precuneus subregions (7A vs.7P), percent signal change, and condition (Own Eyes vs. Retrieve)

**Supplemental Table 25**

*Descriptive statistics in Session 1 and Session 2*

| | Own Eyes | Observer | Retrieve |
|---|---|---|---|
| **Session 1** | | | |
| Memory Age (years) | 9.92 (1.77) | 9.89 (1.89) | 9.88 (1.92) |
| Vividness | 3.65 (.56) | 3.65 (.55) | 3.61 (.54) |
| Emotional Intensity | 3.33 (.54) | 3.28 (.58) | 3.28 (.56) |
| Positive Valence | 3.41 (.38) | 3.35 (.36) | 3.39 (.44) |
| Own Eyes Perspective | 3.77 (.73) | 3.76 (.73) | 3.80 (.72) |
| Observer Perspective | 2.18 (.70) | 2.18 (.71) | 2.15 (.70) |
| **Session 2** | | | |
| Vividness | 3.53 (.61) | 3.49 (.59) | 3.59 (.55) |
| Vividness RT (s) | 1.26 (.25) | 1.26 (.26) | 1.21 (.25) |
| Emotional Intensity | 3.31 (.66) | 3.25 (.70) | 3.33 (.63) |
| Emotional Intensity RT (s) | 1.33 (.31) | 1.34 (.27) | 1.31 (.22) |
| Perspective Maintenance | 3.74 (.69) | 3.42 (.61) | 3.86 (.60) |
| Perspective Maintenance RT (s) | 1.18 (.33) | 1.23 (.33) | 1.24 (.34) |

Note. *Mean* (*SD*)

**Supplemental Table 26**

*Regions Commonly Activated in the Conjunction Analysis for the Cue and Probe Phases (Own Eyes > Retrieve ∩ Observer > Retrieve)*

| | | | | MNI peak | | |
|---|---|---|---|---|---|---|
| Region | Voxels | BA | t | x | y | z |
| **Cue** | | | | | | |
| Visual Cortex | 170 | 19 | 4.54 | -42 | -76 | 22 |
| | | 19 | 3.32 | -38 | -66 | 18 |
| Angular Gyrus | | 39 | 4.08 | -48 | -70 | 18 |
| **Probe** | | | | | | |
| *no suprathreshold clusters* | | | | | | |

MNI = Montreal Neurological Institute; BA = Brodmann's Area;

$p$ = .001, FWEc = 170.

**Supplemental Table 27**

*Results of the Paired Samples T-test for the Cue and Probe Phases*

| | | | | MNI peak | | |
|---|---|---|---|---|---|---|
| Region | Voxels | BA | t | x | y | z |
| **Cue** | | | | | | |
| **Own Eyes > Observer & Retrieve** | | | | | | |
| *no suprathreshold clusters* | | | | | | |
| **Observer > Own Eyes & Retrieve** | | | | | | |
| Visual Cortex | 487 | 18 | 6.52 | 26 | -92 | 2 |
| | | 18 | 6.01 | 26 | -98 | -6 |
| | | 18 | 5.13 | 32 | -94 | 8 |
| | 366 | 18 | 5.57 | -18 | -104 | -6 |
| | | 18 | 5.46 | -26 | -100 | 4 |
| | | 18 | 5.25 | -26 | -100 | -8 |
| **Probe** | | | | | | |
| **Own Eyes > Observer & Retrieve** | | | | | | |
| *no suprathreshold clusters* | | | | | | |
| **Observer > Own Eyes & Retrieve** | | | | | | |
| Posterior Cingulate Cortex | 265 | 31 | 6.58 | -8 | -60 | 44 |
| | | 31 | 4.33 | 0 | -54 | 40 |
| Precuneus | | 7 | 3.59 | 0 | -70 | 50 |
| Anterior Prefrontal Cortex | 281 | 10 | 5.54 | 36 | 52 | -4 |

| | | | | | | |
|---|---|---|---|---|---|---|
| | | 10 | 4.70 | 24 | 48 | 2 |
| | | 10 | 4.30 | 26 | 62 | -6 |
| | 163 | 10 | 5.12 | -32 | 52 | 8 |
| | | 10 | 4.90 | -30 | 52 | -4 |
| | | 10 | 4.30 | -30 | 60 | -2 |
| Angular Gyrus | 116 | 39 | 5.03 | -40 | -80 | 38 |
| Visual Cortex | | 19 | 4.77 | -40 | -84 | 30 |
| | | 19 | 4.35 | -40 | -74 | 26 |

MNI = Montreal Neurological Institute; BA = Brodmann's Area;

Cue: $p$ = .001, FWEc = 366. Probe: $p$ = .001, FWEc = 116.

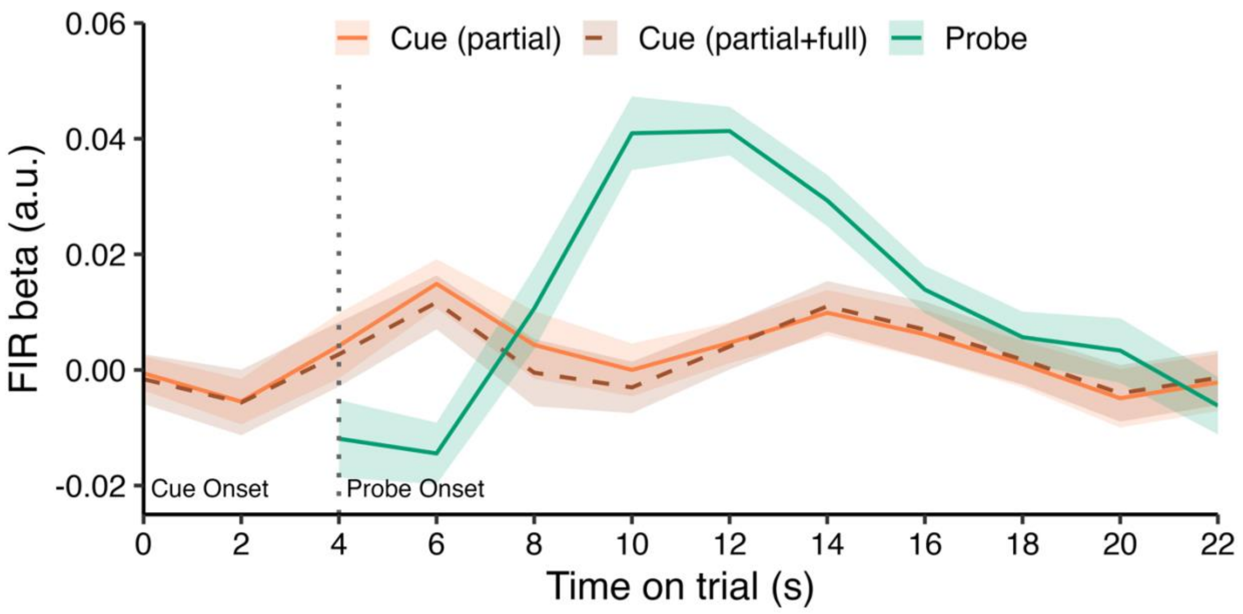


**Supplemental Figure 1.** Finite impulse response (FIR) timecourses (2-s bins, mean ± 95% CI) averaged across conditions (Own-Eyes, Observer, Retrieve) in the left angular gyrus to illustrate temporal separation of cue- and probe-related activity. Cue responses (solid orange line) were estimated from partial trials, showing a distinct early peak at ~6 s. Probe responses (solid teal line, shifted to trial time for display) exhibited a later and larger peak at ~10–12 s. The dashed brown line shows cue-related activity when combining partial and full trials, which closely overlaps with cue-only estimates in the early bins. This demonstrates that including both partial and full trials in the model allows reliable separation of cue- and probe-related responses despite their temporal proximity. The vertical dotted line marks the probe onset (4 s).

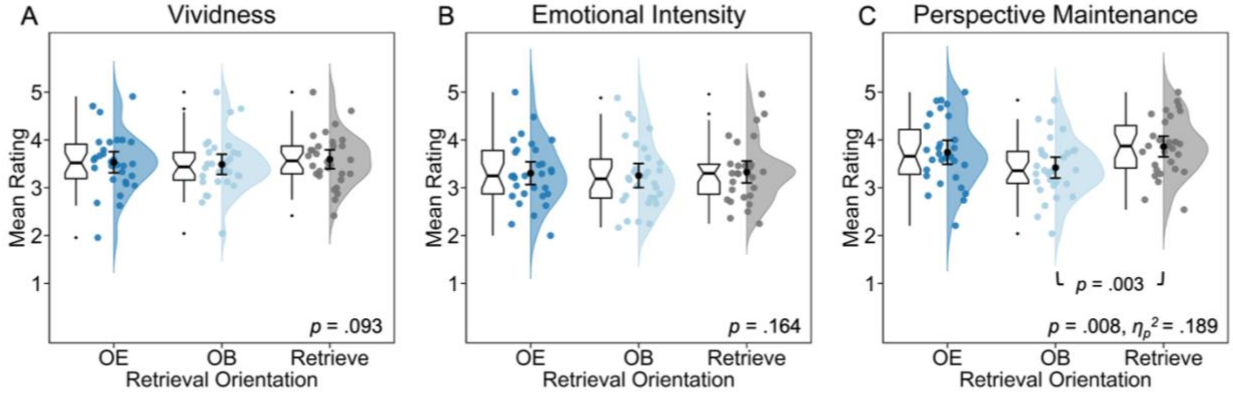


**Supplemental Figure 2.** Online behavioral ratings during fMRI scanning (Session 2), averaged within each participant. Results showed no significant difference for (A) Vividness and (B) Emotional Intensity. However, participants were better able to (C) maintain their natural perspective in the Retrieve condition, compared to remembering AMs from an observer-like perspective.

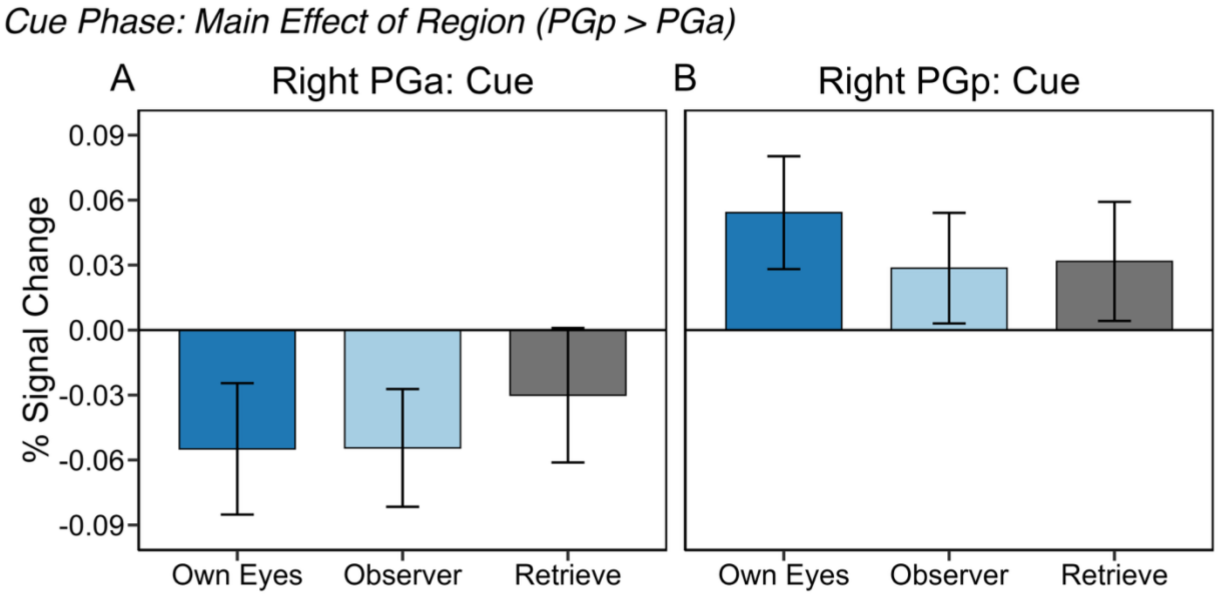


**Supplemental Figure 3**. ROI analysis in right angular gyrus (AG) for PGa (A) and PGp (B).
